# Canonically minimal RNA-guided insertion sequences expand into large elements that disseminate antimicrobial resistance

**DOI:** 10.64898/2026.08.14.744561

**Authors:** Kuang Hu, Bingliang Xie, HengYi Yang, Benjamin E. Rubin

**Affiliations:** Innovative Genomics Institute, University of California, Berkeley, CA 94720, United States

## Abstract

IS110 has emerged as a powerful genome-editing tool because it is the smallest RNA-guided system capable of diverse programmable insertions. Naturally existing elements are conventionally modeled as compact ∼1.5-kb systems comprising a single transposase and a bridge RNA (bRNA). Using high-throughput junction mapping together with large-scale comparative genomics, we redefined the *in vivo* structural boundaries, growth, and mobilization of IS110 elements. We uncovered a previously unrecognized size continuum extending to ∼100 kb, driven by progressive additions. Experiments confirmed activity of natural IS110s both well below and above the size range of previously characterized elements. The large systems comprised mostly plasmid-derived DNA, with antimicrobial resistance genes being the most enriched. Boundary configurations at expanded loci and the range of partial excision intermediates they produce both indicate flexible sequence recognition by IS110, with half-matches between the bRNA and complementary DNA sequence as the most enriched configuration. This sequence tolerance allows loci to expand with diverse cargo. Together, these findings redefine IS110 from a compact insertion sequence into a dynamic platform that disseminates adaptive cargos.

## Introduction

The ability to rewrite genomic DNA at defined loci is foundational to both biological research and therapeutic genome engineering. Most current programmable editing technologies achieve this through targeted DNA cleavage coupled to cellular repair [3, 13, 29]. However, reliance on DNA repair constrains editing efficiency and precision [2]. Double-stranded cleavage is frequently lethal in bacteria, and in eukaryotes repair-dependent strategies lead to inconsistent editing outcomes [15, 32]. The recent discovery of bridge RNA (bRNA)-guided recombinases in the IS110 insertion sequence family offers an alternative that both finds the target site and inserts the specified cargo directly, rather than generating a break that depends on host repair [18]. Unlike CRISPR-associated transposase (CAST) systems, which require multiple protein components for targeted insertion [31, 61], IS110 recombinases operate through a single RNA-guided protein [25]. The bRNA contains independently programmable donor- and target-recognition sites that can be reconfigured to specify new sites and inserted cargoes [18]. Together with the recombinase, this architecture enables targeted insertions, inversions, and excisions without a double-stranded break or homology-directed repair [25, 55]. Engineering studies have demonstrated efficient large-fragment insertion and megabase-scale genome rearrangements, highlighting the potential of IS110 as a programmable genome engineering platform [45, 47]. Furthermore recent biochemical and structural analyses have began to resolve how these systems mobilize in nature [68]. Yet these mechanistic and engineering studies remain centered on a small number of experimentally tractable systems, leaving the structural diversity, mobility, and evolutionary behavior of IS110 across natural genomes largely unexplored.

Previously, IS110 elements have been defined as compact, ∼1.5-kb RNA-guided systems comprising a single recombinase-coding sequence and a short bridge RNA (bRNA) [11, 18, 25]. This fits the broader, conventional definition of insertion sequences as mobile elements under 2.5 kb that encode only the functions required for their own transposition [38]. However, this model—inferred from a limited set of experimentally characterized systems and pipelines optimized for canonical insertion sequences—may systematically overlook extended non-coding regions and alternative structural organizations [56, 57, 69]. Whether the canonical 1.5-kb model accurately captures the full structural and functional diversity of the IS110 family therefore remains unknown.

To determine the true footprint of active IS110 elements and assess their diversity across natural genomes, we developed a framework combining high-throughput read-to-reference junction mapping, circular intermediate analysis, and large-scale comparative genomics. Using this approach, we found that IS110 spans a previously unrecognized size continuum, from elements only ∼67% the size of the smallest previously validated IS110 (800 bp versus 1,202 bp) to loci approaching 100 kb, nearly 40-fold larger than the largest previously characterized WT IS110 (2,527 bp). Evidence for whole-unit mobility, however, was largely restricted to elements under 20 kb. We further experimentally validated the activity of IS110 systems both significantly smaller and larger than those previously characterized, spanning from an 843-bp minimal element to a 9.2-kb expanded locus. The expanded IS110s preferentially accumulate adaptive cargo, including antimicrobial resistance determinants and heavy-metal detoxification systems, are enriched for plasmid-derived DNA, and participate in extensive cross-genus horizontal gene transfer, revealing IS110 as an important vehicle for the dissemination of adaptive genetic material. Analysis of excision intermediates and expanded loci showed that their boundaries retain only a single identifiable canonical donor-boundary match, revealing underappreciated flexibility in recombinase substrates. Together, these findings redefine IS110 as an often large mobile element mobilizing adaptive cargo across genera.

## Results

### Identification and Diversity of Large Cargo-containing IS110 Elements

To define the *in vivo* boundaries and activity of IS110 elements, we combined read-to-reference comparisons with circular-junction analysis (Fig. 1A, Fig. S1). We first identified likely mobile genetic element boundaries by comparing occupied genomic loci with the corresponding empty reference sites [35, 59, 65]. Once these boundaries were established, we searched for long reads spanning the joined ends of the element to detect circular recombination intermediates, providing direct evidence that the reconstructed element undergoes mobilization. We then filtered to those bounded sequences that contain a gene with IS110 domains, enabling large-scale characterization of canonical and expanded IS110 architectures. To determine whether expanded loci represented *bona fide* IS110-associated elements rather than unrelated mobile elements carrying an IS110 recombinase, we examined their coding content and predicted bRNA architecture (Fig. S2). Among 2,999 elements *>* 2 kb, 1,474 cargo-bearing loci lacked any detectable alternative transposase or integrase (Fig. S2A–B), while LTG/RTG sequences complementary to the corresponding target flanks were enriched in non-coding, bRNA-like regions relative to shuffled controls (Fig. S2C). Together, these features support the expanded loci as IS110-associated structures rather than incidental IS110 passengers within other mobile elements.

**Figure 1.**
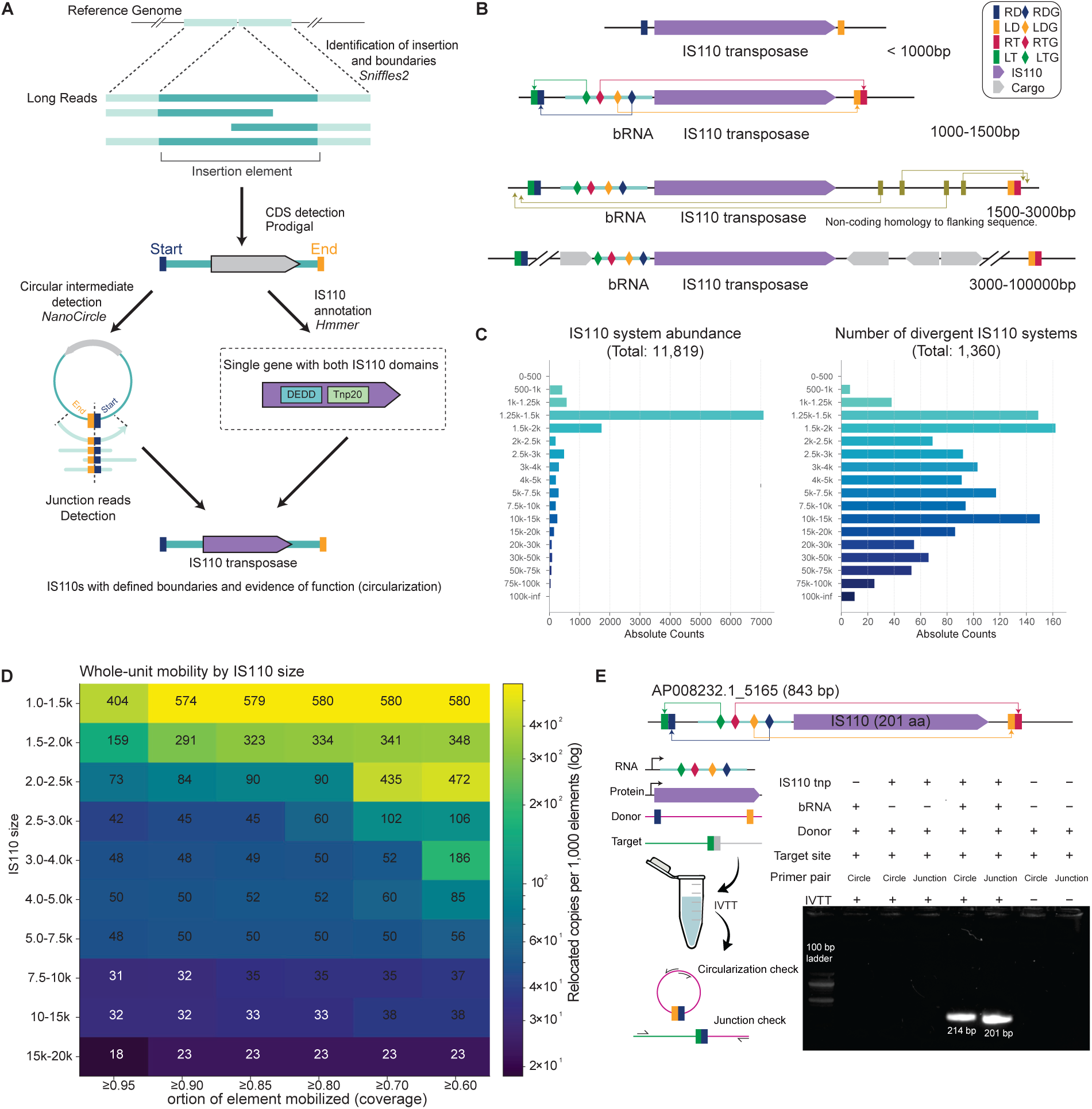
Genome-wide discovery reveals the structural diversity and mobility landscape of IS110 elements. **A.** Overview of the computational framework used to identify functional IS110 elements from long-read sequencing and comparative genomics. Structural variant detection, coding-sequence annotation, and circular-intermediate analysis were integrated to reconstruct complete IS110 boundaries and identify active elements supported by circularization evidence. **B.** Representative architectures of naturally occurring IS110 elements, including minimal transposase-only elements, canonical bRNA-containing systems, elements with extended non-coding regions, and expanded cargo-bearing loci approaching 100 kb. **C.**Length distribution of Nanopore-supported IS110 loci before (11,819 loci) and after dereplication at 95% nucleotide identity (1,360 representatives), revealing a continuous size spectrum extending from canonical ∼1.5-kb elements to nearly 100 kb. **D.** Normalized recovery of identical IS110 bodies across distinct genomic contexts as a function of element length and alignment coverage threshold. **E.** Cell-free validation of a compact IS110 element (843 bp, 201 aa). An IVTT one-pot assay reconstituting the bRNA, IS110 transposase, donor, and target substrates produced PCR-detectable donor circularization and integration junctions only in the presence of both the transposase and bRNA.

Contrary to the prevailing view that IS110 consists of a compact recombinase–bridge RNA module [11, 18], our analysis revealed a substantially broader spectrum of genomic architectures (Fig. 1B). In addition to canonical elements, we identified variants containing extended 3*^/^* non-coding regions with sequence homology to flanking chromosomal DNA, as well as numerous loci carrying multi-gene cargo. These architectural differences were accompanied by an unexpectedly broad size distribution among long-read-supported IS110 loci. Although canonical IS110 elements clustered tightly between approximately 1.0 and 1.6 kb, the diversity of systems (redundancy reduction at 95% nucleotide identity) had a continuous distribution extending to ∼100 kb, with a dereplicated median size of approximately 4 kb—roughly twice the size of the largest previously characterized IS110 element (Fig. 1C). We also recovered a circular molecule joining the ends of a 9,177-bp IS110 locus by rolling-circle amplification and long-read sequencing, indicating that native IS110 recombination mobilize regions far larger than previously appreciated (Fig. 5B). Consistent with the ability of the assay to discriminate functional IS110s, elements carrying truncated recombinases showed no evidence of activity. Rather than representing rare outliers, expanded IS110 loci were widespread across our dataset, indicating that the canonical ∼1.5-kb architecture captures only a subset of the natural diversity of the IS110 family.

Despite this remarkable size expansion, direct evidence for whole-unit mobility showed the opposite trend. Circular intermediates and relocation of identical element bodies between distinct genomic contexts were most frequently detected among canonical 1.0–1.6 kb elements and declined progressively with increasing size. In these larger systems, most mobilization was of partial segments. Above approximately 20 kb, we detected no evidence of whole-unit mobilization. Instead, larger loci continued to expand through cargo acquisition and tandem duplication, indicating that size expansion and whole-unit mobility become progressively decoupled during IS110 evolution (Fig. 1D).

We also identified many IS110s smaller than those previously characterized. These include rare, putatively active elements of approximately 800 bp that retain an intact recombinase coding sequence despite lacking the 5*^/^* untranslated region, suggesting that bridge RNA can function *in trans*. Also in this set are IS110 elements complete with putative bridge RNAs that are far more compact than those previously tested. We experimentally validated the activity of an exceptionally compact 843-bp IS110 element encoding a 201-amino-acid recombinase by reconstituting the system in a cell-free *in vitro* transcription–translation (IVTT) assay, confirming both donor circularization and target integration by PCR and Sanger sequencing (Fig. 1E, Fig. S3). Its predicted guide–DNA matches were also relatively long (LTG, 10 bp; RTG, 9 bp; LDG, 9 bp; RDG, 8 bp with one mismatch). In comparison, the canonical IS110-family element adapted for editing is the 1,279-bp IS110 IS621 encoding a 326-amino-acid recombinase [11].

### IS110 Elements are Enriched for Adaptive Cargo

We sought to determine whether the expanded regions of IS110 loci are similar in content to their host genome or selectively retain certain types of cargo. To do this, we compared the functional composition of coding sequences from cargo-bearing IS110 elements with those of plasmids and length-matched genomic fragments [53] (Fig. 2A,B). Both IS110 and plasmids were depleted in core metabolic functions relative to the genomic background (∼ 0.4× and ∼ 0.2×, respectively) [9, 26], consistent with mobile genetic elements acting primarily as vehicles for adaptive rather than housekeeping functions [50]. Conversely, antimicrobial resistance (AMR) emerged as the most overrepresented feature of IS110 cargo [1, 6, 20]. AMR determinants were enriched approximately 30-fold over the genomic background and twice that observed for plasmids, accounting for approximately 13% of all non-backbone CDSs. Defense genes were also enriched in IS110, although to a much lesser extent than in plasmids (∼ 2× versus 9× in plasmids) [63]. At the subclass level, resistance-associated IS110 cargo was dominated by sulfonamide, aminoglycoside, and *β*-lactam resistance together with Zn/Cd/Co/Ni and mercury detoxification (Fig. 2C). Metabolic subclasses remained abundant within IS110 cargo in absolute terms, particularly amino acid, energy, and ion-associated functions, but were depleted relative to the genomic background (Fig. 2B,C). Therefore, IS110 cargo does not mirror the host genome. Its pattern of gene depletion and enrichment broadly follows plasmids, but with divergences such as the enrichment of AMR cargo.

**Figure 2.**
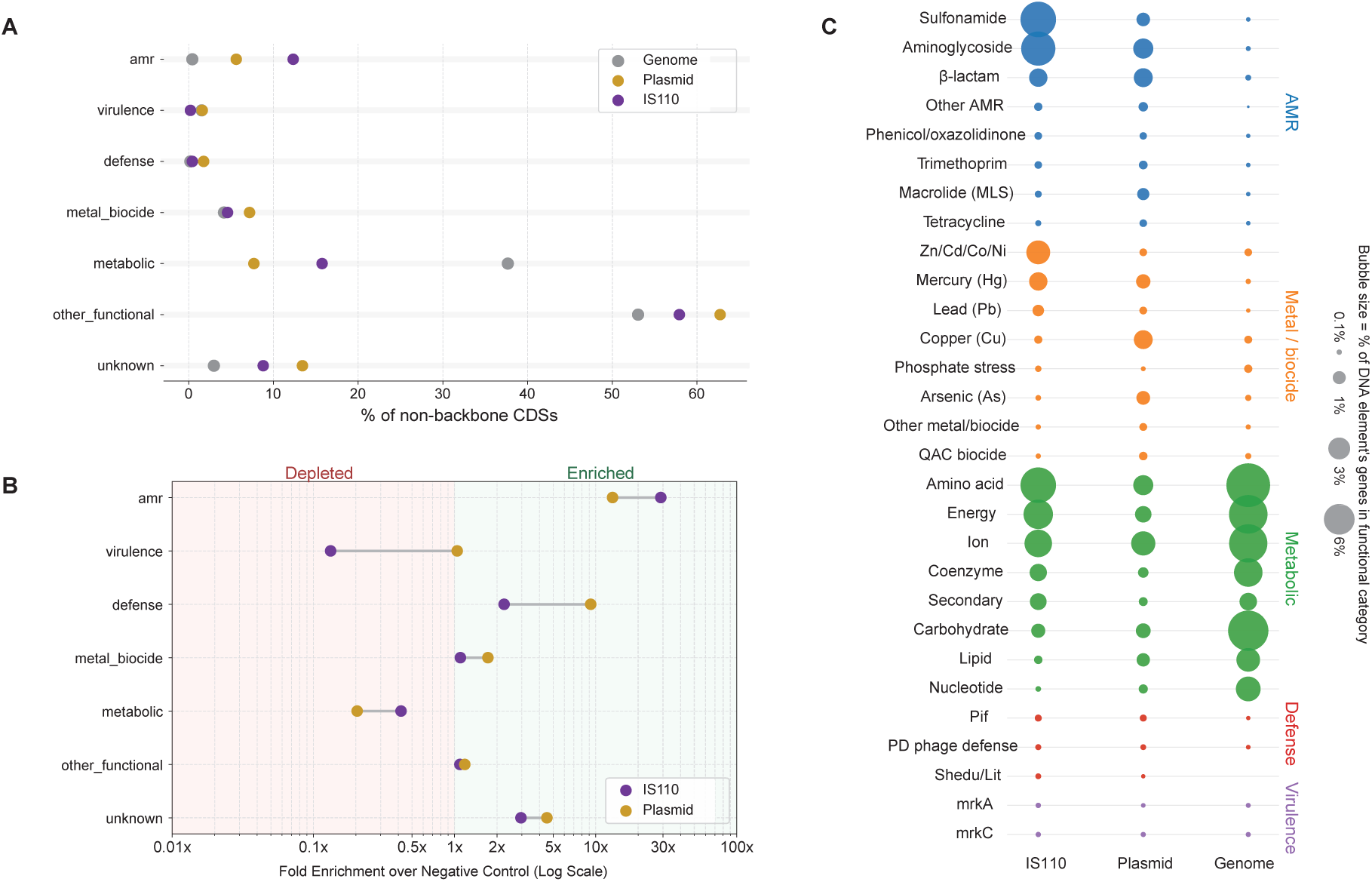
Expanded IS110 loci preferentially accumulate adaptive cargo. **A.** Relative abundance of major functional categories among non-backbone coding sequences from IS110 loci, plasmids (mobile genetic element control), and length-matched genomic fragments (negative control). **B.** Fold enrichment of each functional category in IS110 loci and plasmids relative to genomic controls. Antimicrobial resistance genes showed the strongest enrichment in IS110 cargo, whereas core metabolic functions were depleted in both mobile genetic element datasets. **C.** Functional subclass composition of adaptive cargo carried by IS110 loci, plasmids, and genomic controls. Bubble size represents the proportion of genes assigned to each functional subclass within a dataset.

### Large Plasmid-like IS110s Transfer Adaptive Cargo Across Genera

To investigate whether expanded IS110 loci transfer across bacterial lineages, we searched representative IS110 loci ≥5 kb (*n* = 1,244 after 99% dereplication) against complete genome collections from nine bacterial genera. Candidate transfer events were required to share highly conserved IS110 sequences (≥99% nucleotide identity and ≥90% coverage) while exhibiting minimal conservation of the surrounding chromosomal context (*<* 10% coverage and *<* 50% similarity), thereby distinguishing recent horizontal transfer from shared ancestry [58]. Although these criteria may exclude instances where the RNA-guided targeting of IS110s leads to integration into homologous sites across different genera, this conservative threshold was intentionally chosen to minimize false positives. Using these stringent criteria, we identified approximately 30,000 cross-genus transfer events representing 615 distinct IS110 variants (Fig. 3A). Transfer networks were particularly dense among *Klebsiella*, *Enterobacter*, *Escherichia*, and *Salmonella*, demonstrating that large IS110 loci undergo widespread exchange among clinically important *Enterobacteriaceae*.

**Figure 3.**
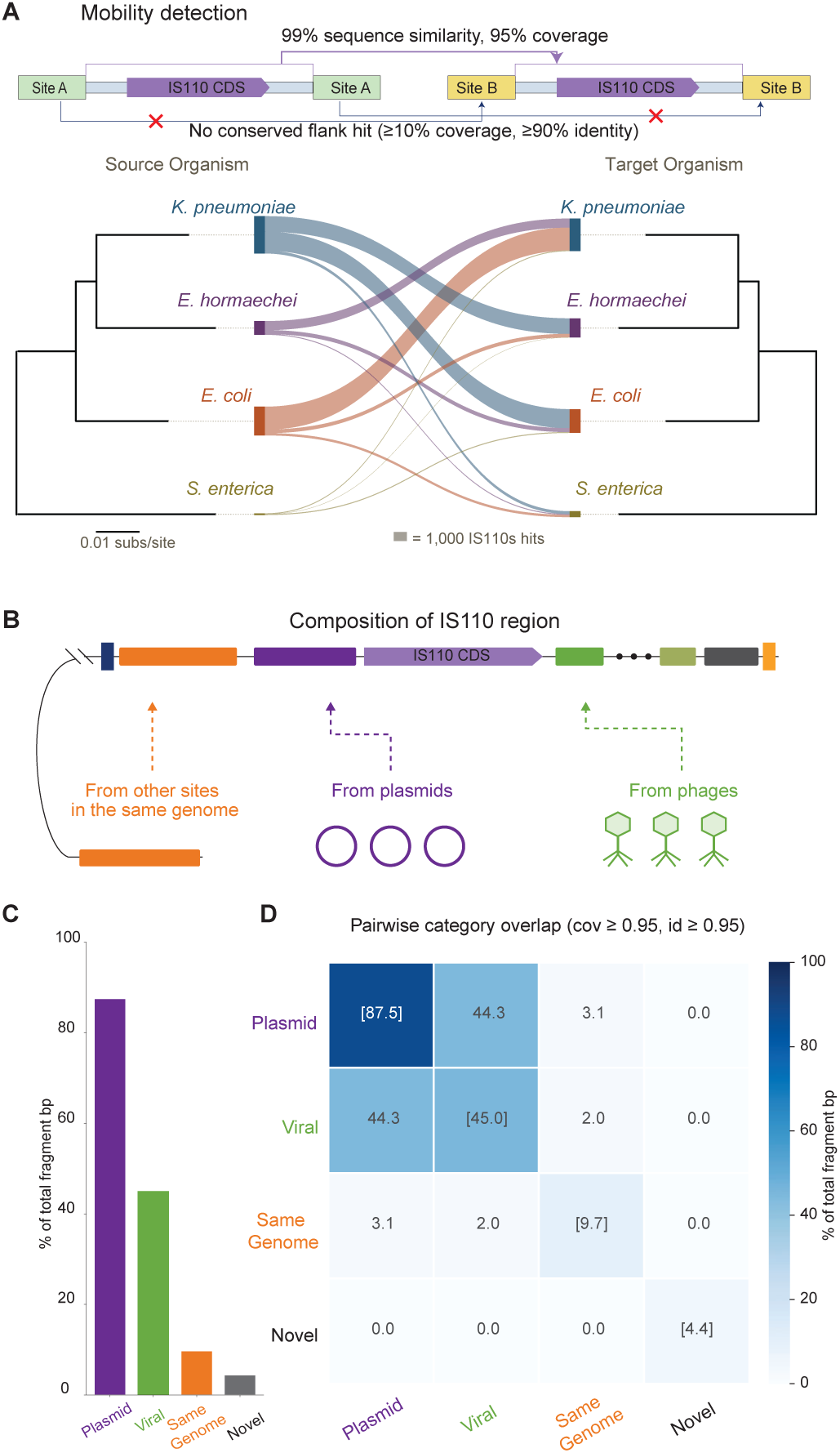
Expanded IS110 loci undergo extensive horizontal transfer and accumulate mobile genetic element-derived DNA. **A.** Cross-genus transfer analysis of representative IS110 loci (≥5 kb) identified highly conserved IS110 bodies (≥99% nucleotide identity and ≥95% coverage) embedded within divergent chromosomal contexts (*<* 10% coverage and *<* 90% sequence similarity of the flanking regions). The ribbon plot summarizes high-confidence IS110 matches between source (left) and target (right) species; ribbon width is proportional to the number of pairwise matches, and the flanking dendrograms show the phylogenetic relationships of the four species with the densest transfer networks (scale bar, 0.01 substitutions per site). **B.** Schematic illustrating the inferred origins of DNA incorporated into expanded IS110 loci, including chromosomal DNA from other genomic locations, plasmids, bacteriophages, and previously unclassified sequences. **C.** Percentage of IS110 body sequence matching each inferred source category (coverage ≥ 95%, identity ≥ 95%; equivalent to the diagonal of panel D). Categories are non-exclusive and may sum to *>* 100% because individual sequences can match multiple source categories. Most IS110 body sequence matched plasmid references, with substantial additional matching to phage references, whereas comparatively little matched other chromosomal regions. **D.** Pairwise overlap among inferred source categories based on high-confidence sequence matches (coverage ≥95%, identity ≥95%), revealing substantial sequence sharing between plasmid- and phage-derived regions.

To determine the origin of the DNA accumulated within expanded IS110 loci, we classified their sequences using complementary MOB-suite and geNomad annotations benchmarked against random genomic controls [8, 52]. The vast majority was from mobile genetic elements, particularly plasmids (Fig. 3B,C). Chromosomal sequences also represented a minor, but detectable share. Pairwise overlap analysis further revealed substantial sequence sharing between plasmid- and phage-derived annotations (Fig. 3D), indicating that many captured regions represent shared elements rather than DNA unique to either source. These findings establish expanded IS110 loci as hubs for the acquisition and exchange of mobile genetic material.

The extensive contribution of plasmid-derived DNA to expanded IS110 loci prompted us to ask whether intact IS110 elements themselves also shuttle between chromosomes and plasmids. Comparison of chromosomal IS110 loci with the PLSDB plasmid database identified 305 high-confidence chromosome–plasmid matches, representing 12% of all dereplicated IS110 lineages (Fig. S4A,B). Shared IS110 elements typically occupied a fraction of their host plasmids (Fig. S4C), indicating that chromosome–plasmid shuttling generally involves IS110 embedded within larger plasmid backbones rather than transfer of entire plasmids. A few expanded loci carry this to its limit, retaining plasmid replication and transfer determinants (Fig. S4D) that leave little to distinguish the circular intermediate from a small plasmid.

### Repeated Insertions and Duplications Generate Expanded IS110 Loci

To determine how giant IS110 loci arise, we compared closely related IS110 insertions occupying the same chromosomal location across related genomes. We identified numerous examples in which additional DNA accumulated progressively within an otherwise conserved insertion site while the surrounding chromosomal context remained unchanged (Fig. 4A–C). These observations support stepwise *in situ* growth as a mechanism contributing to the formation of expanded IS110 loci. Beyond simple insertional growth, expanded loci also showed recurrent tandem duplication of full or partial IS110 modules (Fig. 4B,C). Expanded IS110 loci were also found to undergo inversions, inter-chromosomal translocations, and tandem duplications (Fig. S5). Recurrent insertion into already-occupied regions also implies that IS110 lacks an equivalent of the target immunity that suppresses insertion near existing copies in the Tn3 and Tn7 families [4, 34, 60]. Such a mechanism may be dispensable for IS110, since insertion splits the target into LT|RD and LD|RT hybrid junctions and does not regenerate an intact target site [18, 62]. Together, these features support a model where repeated local insertion and duplication drive the progressive expansion of IS110 loci.

**Figure 4.**
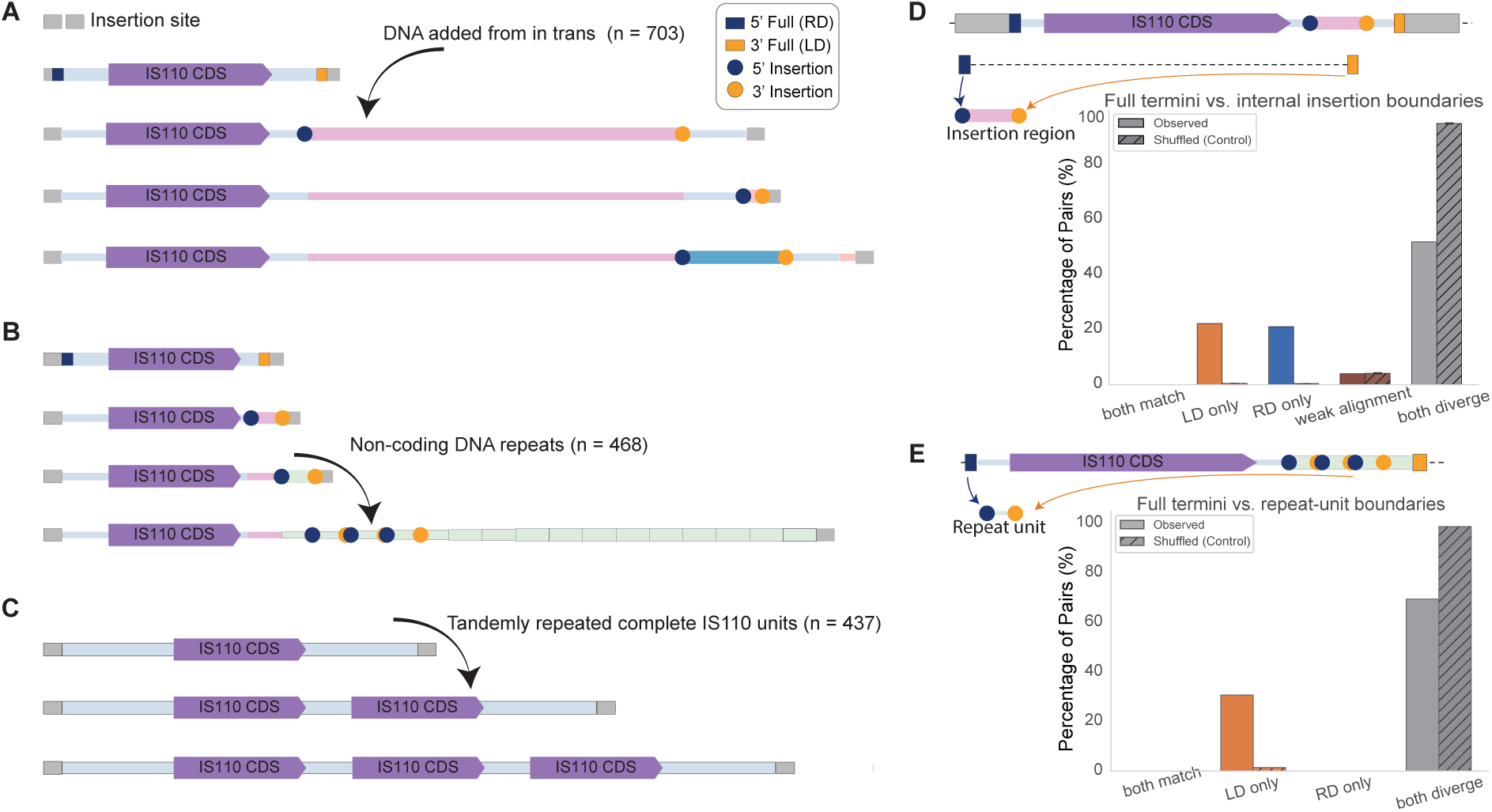
Local expansion of IS110 loci proceeds through sequential insertion and tandem duplication. **A.** Representative examples of progressive expansion through sequential acquisition of DNA in trans downstream of a conserved IS110 transposase module (n = 703). **B.** Representative expansion events involving tandem accumulation of non-coding repeat units adjacent to the IS110 transposase (n = 468). **C.** Representative tandem duplication of complete IS110 modules, generating multi-copy composite loci (n = 437). **D.** Comparison of donor-boundary matching frequencies between complete IS110 termini and internal insertion boundaries. Left-donor (LD)-only and right-donor (RD)-only matches were enriched at authentic insertion boundaries relative to shuffled controls. **E.** Comparison of donor-boundary matching frequencies at repeat-unit boundaries generated during tandem expansion. Repeat-unit boundaries preferentially retained LD-only matches, consistent with progressive local expansion initiated from authentic IS110 donor boundaries.

To determine whether local expansion bears signatures of IS110-specific recombination rather than generic recombination, we compared the boundary sequences of full IS110 elements with those of internal insertions and repeat units (Fig. 4D,E). We reasoned that if local expansion repeatedly engages the resident IS110 recombination system, expansion-associated boundaries should retain donor sequences recognizable by the same bRNA guide architecture. Relative to shuffled controls, internal insertion boundaries were strongly enriched for sequences matching the donor ends of the element that housed them (LD and RD). Interestingly, these matches were almost always one-sided. Individual boundaries carried either an LD or an RD match, while complete donor pairs were essentially absent (Fig. 4D,E). This aligns with recent biochemical evidence that IS621 recombination can initiate at half-matches to the bRNA [68]. In contrast to canonical recombination, where the inserted product replaces sequence between the terminal donor sequences, such a half match could support element expansion. Although perfect matches to the host IS110’s likely LD and RD were detectable at only a subset of boundaries, their enrichment relative to shuffled controls indicates that expansion recurrently engages the resident IS110 recombination system, with the sequences recognized at the remaining boundaries yet to be defined.

### Expanded IS110 form partial excision-intermediates with flexible terminal sequences

Once an IS110 element has been inserted, its two donor half-sites are separated between the hybrid junctions LT|RD and LD|RT, neither of which provides a full match to the bRNA for element excision and formation of its circular intermediate. Recent work shows that the full donor match required for excision can be reconstituted by bringing the element’s two hybrid junctions into proximity [24], or bypassed altogether when initiation occurs from LD- and RD-half-matches [24, 68]. In addition, modular circularization has been observed in ISPpu9, where alternative combinations of related boundary sequences generate distinct sub-element circular excision products [44]. We therefore examined circular intermediates across expanded IS110 loci to determine whether partial circularization is a broader feature of natural IS110 systems and what sequence features define these alternative boundaries.

Across our dataset, alongside full-length circles, we recovered numerous circular intermediates strictly shorter than their parent elements, hereafter referred to as partial circles, to differentiate them from the ISPpu9 ‘minicircles’ that encompass full intermediates. These products were particularly prominent at expanded loci. For example, an 18.4-kb locus produced two distinct partial circles spanning 19% and 79% of the element, with no detectable whole-element circle, whereas a canonical 1.3-kb element produced abundant full-length circles (Fig. S6). Partial circularization thus becomes increasingly prominent across the size range in which whole-unit mobility declines (Fig. 1D).

To determine which donor boundaries support these products, we compared partial-circle junctions with the full-length circle junction from the same element (Fig. 5A). Similar to the insertion boundaries (Fig. 4), excision partial circles rarely retained matches to both canonical donor boundaries that are expected to be recognized by the bRNA donor loop. Instead, LD-only and RD-only configurations were enriched relative to shuffled junction controls, indicating that many partial circles retain only a single assignable canonical guide–donor match. For LD-only and RD-only junctions, the unmatched boundary showed no greater similarity to the corresponding canonical donor sequence than expected by chance (Fig. S7). Junctions classified as divergent at both boundaries nevertheless constituted the largest observed class in absolute terms, although they were substantially depleted relative to shuffled controls. Taken together, the enrichment of single-match configurations and the absence of residual similarity at the unmatched boundary indicate that partial circularization is frequently guided by a single half-match to the parent IS110’s predicted LD or RD, rather than requiring both.

**Figure 5.**
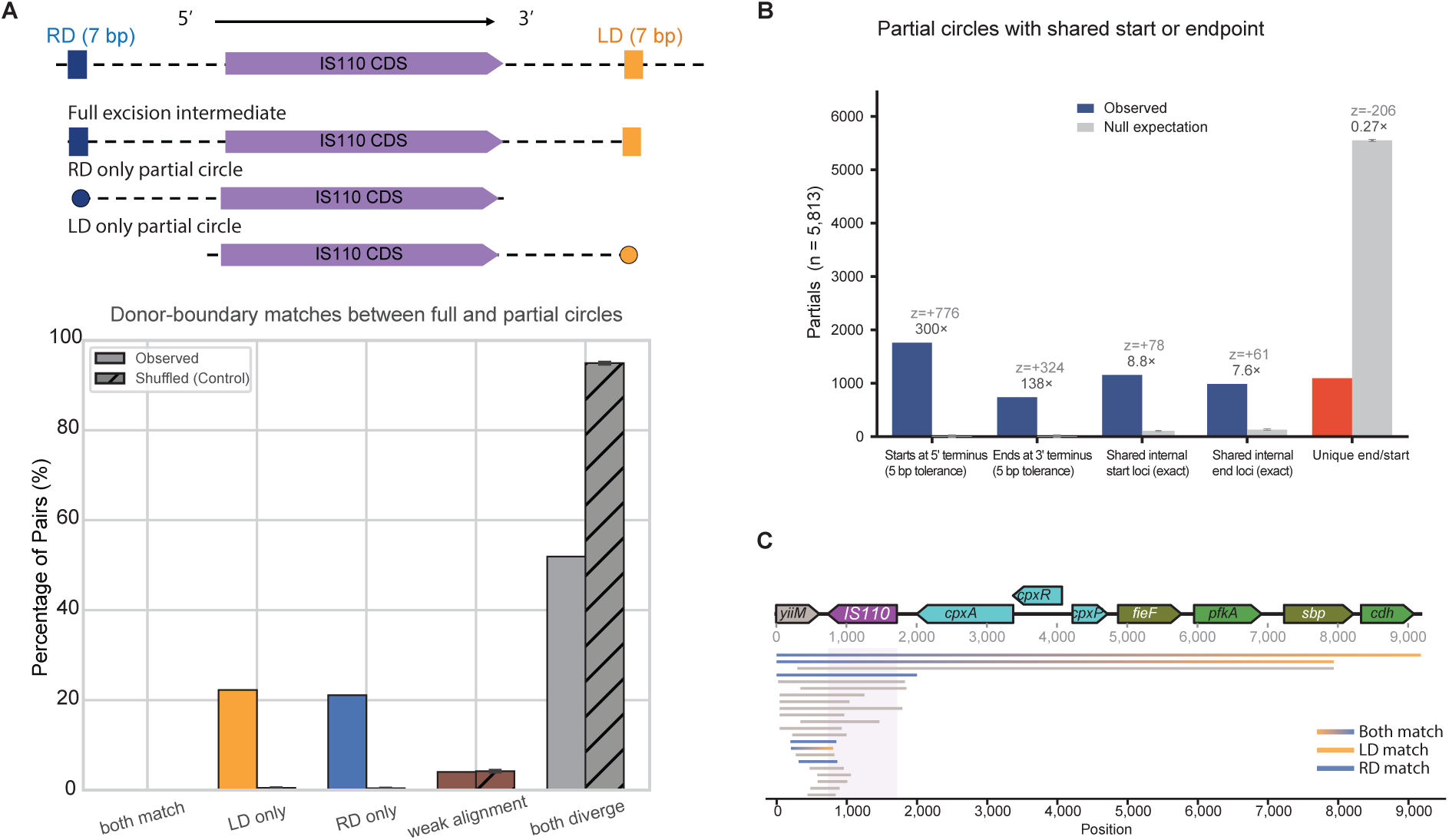
Partial IS110 circularization frequently retains only one canonical donor-boundary match. **A.** Junction-side sequence comparison between full-length and partial circular intermediates. The schematic illustrates the regions compared at the circular junction, corresponding to the right-donor (RD; 7 bp) and left-donor (LD; 7 bp) guide-matching sequences. For each full–partial circle pair, the two junction sides were classified according to whether the corresponding LD and RD regions matched (both match, ld only, rd only, weak alignment, or both diverge). The bar plot shows the frequency of each class in observed full–partial pairs relative to shuffled controls. **B.** Bar plot of partial-circle endpoint sharing relative to full IS110 termini and other partial circles. Observed endpoint-sharing counts among 5,813 partial circles were compared with a randomized null expectation. Red denotes the observed “Unique end/start” category, which is depleted relative to the randomized null. Full-element endpoint reuse was scored within a 5-bp tolerance, whereas partial–partial endpoint sharing required an exact positional match. **C.** Rolling-circle amplification (RCA) reads mapped to reconstructed circular IS110 products. Full-length and partial circular intermediates from the same locus were recovered as discrete circular products that differed in the extent of retained sequence, independently supporting the partial-circle intermediates detected by long-read junction analysis.

Two further features of the partial-circle coordinates indicate that their boundaries are strongly non-random (Fig. 5B). First, many partial circles reused a terminal boundary of the full element: reuse of the left and right termini was enriched 300-fold and 138-fold, respectively, relative to randomized endpoints. Second, internal endpoints recurred across independent partial circles, with shared internal start and end positions enriched 8.8-fold and 7.6-fold, respectively, whereas partial circles sharing no endpoint with either the full element or another partial circle were depleted to 0.27-fold of the null expectation. Partial-circle termini are therefore not determined solely by canonical LD and RD homology, but are concentrated at a restricted set of recurrent positions within IS110 loci.

At least some divergent termini sequences in partial sequences could be explained by alternative guide assignments. In one case, a nearby 6–7-nt sequence within the same exposed bRNA loop provided an alternative donor-recognition segment compatible with the observed junction sequence (Fig. S8). In another, a single IS110 locus contained multiple spatially distinct predicted donor-binding loops associated with different partial-circle junctions (Fig. S9). In these cases, the donor sequence used appears to shift within a predicted bRNA loop or between loops, so a single IS110 element can support more than one donor-binding configuration. Because our classification used stringent alignment criteria for these short recognition segments, genuine but imperfect guide–donor pairings may therefore be underestimated, contributing to the relatively low apparent match rate.

Partial IS110 intermediates also showed evidence of genomic reintegration. Comparative genomic analysis identified highly similar (*>* 99% nucleotide identity) partial IS110 fragments inserted at independent chromosomal sites, together with the corresponding empty donor and target loci (Fig. S10). These observations support the ability of partial IS110-derived fragments to reintegrate into bacterial genomes rather than representing sequencing or circle-detection artifacts.

To determine whether these partial products occur in cells, we profiled circular DNA from an *E. coli* strain carrying a naturally occurring large IS110 locus. Circular DNA was enriched from cells carrying the construct using rolling-circle amplification and then sequenced. Full-length and partial circles were recovered (Fig. 5C), experimentally confirming the range of circularization products that can occur from a naturally occurring large IS110 locus. Importantly, these products included junction configurations retaining only one canonical donor-boundary match, experimentally supporting partial-circle formation from boundaries that diverge from the canonical LD–core–RD configuration. Also, consistent with the reanalysis of previous sequencing data (Fig. 5B), the partial circles produced experimentally here (Fig. 5C) cluster around specific start points, end points, or both, indicating non-random formation despite the absence of conventional matches to the bRNA loops. Together, the natural and experimental intermediates show that partial circularization is common in large IS110s and proceeds without matches to both canonical donor boundaries.

## Discussion

Current models define IS110 as a compact ∼1.5-kb RNA-guided insertion sequence. Our analyses instead reveal that natural IS110 systems span a continuum from ∼800 bp to 100 kb. The large IS110s represent evolutionary products of repeated local expansion in which whole-unit mobility becomes progressively uncoupled from locus growth. That cargo comes largely from plasmids and is biased toward adaptive functions, most prominently antimicrobial resistance. This positions acquisition of new DNA elements, rather than transposition alone, as an important mode of IS110 evolution.

The long-standing view of IS110 as a compact insertion sequence likely reflects how these elements have been annotated. Conventional insertion sequence annotation pipelines are optimized to identify transposase genes and nearby terminal features [18, 48, 69], and cannot reconstruct scarlessly inserted loci that carry extensive non-coding regions and accessory cargo. Our results suggest that expanded IS110 loci have been systematically underappreciated rather than exceptionally rare.

One question raised by this work is why IS110 would favor integration of plasmid DNA. The natural IS110 donor is a double-stranded circle [18, 46], and integration favors double-stranded over single-stranded donors [62]. Plasmids already exist in that form, potentially making them a preferred source of captured DNA. Half-match tolerance at donor boundaries (Figs. 4, 5) [68] and flexible bRNA guide usage (Figs. S9, S10) would broaden which plasmids and plasmid segments are eligible. In exceptional cases the expanded loci may retain elements necessary to operate as a mobile replicating plasmid (Fig. S4D). This raises the possibility that some IS110 circular intermediates acquire the ability to replicate and transfer as plasmids in addition to inserting by RNA-guided recombination.

Recognizing putative donor sequences via a single half-match in expanded IS110s likely serves two purposes: relaxing sequence specificity to allow for more flexible expansions and partial excisions, and driving expansion reactions instead of canonical IS110 replacement reactions. Native half-match excision has recently been shown to occur by a single-strand copy-out that leaves the donor locus intact [62, 68], so in IS110 expansion, such half-matches on a single side plausibly add DNA at a boundary rather than the replacement of sequence between two boundaries during canonical insertion [18, 25]. In line with this model, our results show expanded IS110s usually have only one assignable half-matched donor arm. Furthermore, our data suggest that the two arms do not behave identically. Partial circles preferentially reuse the RD-side terminus of the parent element (Fig. 5B,C), whereas tandem repeat units retain almost exclusively LD matches (Fig. 4E). We speculate that reactions initiating at RD tend to complete circularization and excise (Fig. 5B,C), while those initiating at LD are more often resolved in place, duplicating the intervening segment (Fig. 4E). Ordered strand transfer offers a potential basis for this asymmetry because the RD-containing junction appears to be formed first [62]. Functionally asymmetric ends that channel one-sided reactions toward adjacent-DNA capture and tandem duplication have been described for other insertion sequences, most directly in IS91 [40]. How this asymmetry arises in IS110 is unresolved and a target for future biochemical work.

Beyond their ecological significance, these findings have implications for programmable IS110-based genome engineering. The elements described here expand the pool of candidate scaffolds well past the narrow size range from which current editors derive, and the recovery of circular intermediates and cross-genus transfer events marks which of them are likely active. Among them is an 843-bp element encoding a 201-amino-acid recombinase, the most compact IS110 system shown to catalyze targeted insertion (Fig. 1E), roughly a third smaller than IS621 in both element and protein length, while retaining relatively long predicted guide–DNA complementarity, a key feature for accuracy [11]. Systems this small may relax the delivery constraints that limit larger editors, especially for human delivery [67]. Because IS110s were identified here by evidence of mobilization rather than by sequence similarity alone, they represent a set of candidate editors already filtered for activity.

Collectively, our findings redefine IS110 from a compact RNA-guided insertion sequence into a dynamic evolutionary platform for bacterial genome remodeling. Rather than acting solely as self-propagating mobile elements, IS110-seeded loci can progressively expand, acquire adaptive cargo, exchange DNA among chromosomes, plasmids, and phages, and reshape chromosome architecture. More broadly, our results suggest that RNA-guided recombination can contribute to long-term genome innovation through progressive structural diversification, extending the evolutionary roles attributed to the IS110 family beyond transposition alone.

## Methods

### Data Acquisition, Reference Matching, and Read Alignment

Public long-read sequencing datasets generated using Oxford Nanopore Technologies (ONT) or PacBio platforms were retrieved from the European Nucleotide Archive (ENA) and the Sequence Read Archive (SRA) using Kingfisher [66]. Sample metadata, including BioProject identifiers, organism taxonomy, sequencing platform, and sequencing throughput, were obtained from the SRA metadata service using Biopython [12]. Corresponding reference genomes were downloaded with the NCBI Datasets command-line tool [42], preferentially selecting complete RefSeq assemblies matched to the corresponding organism and strain to maximize alignment accuracy and facilitate precise insertion-site reconstruction.

Long reads were aligned to their matched reference genomes using minimap2 [36] with the `-x map-ont` preset for ONT datasets and the `-x map-pb` preset for PacBio datasets. Alignments were generated with the `-a --MD` options to retain SAM-format alignments together with per-base mismatch annotations and supplementary alignments. Resulting alignments were coordinate-sorted and indexed using samtools [16] and HTSlib [5]. To prevent excessive computational resource consumption from problematic datasets, individual alignment jobs were terminated if they exceeded a wall-clock runtime of two hours.

### Structural Variant Calling and Full Circular Intermediate Detection

Insertion-type structural variants (≥ 100 bp) were identified from coordinate-sorted BAM files using Sniffles2 (v2.7.2) [59] with the parameters `--minsupport 3`, `--minsvlen 100`, and `--allow-overwrite`. For each insertion, the genomic coordinates, consensus insertion sequence reconstructed from supporting reads, and supporting read count were extracted from the Sniffles2 output and recorded for downstream analyses.

To identify full circular intermediates generated by IS110 excision, a tandem [IS][IS] concatemer was computationally constructed by duplicating the consensus insertion sequence end-to-end, thereby creating an artificial head-to-tail junction corresponding to the expected circularization product. All long reads were re-aligned against this concatemer using minimap2 [36] with the `-x map-ont` preset. Junction-spanning reads were identified using a custom Python module (`cycle.circle detect`) implemented with pysam (v0.23.3) [49].

Reads containing two alignment segments that both mapped within the duplicated IS sequence and collectively spanned the synthetic head-to-tail junction, with at least 100 bp of aligned sequence on each side of the junction, were classified as *tail-head* (TH) junction reads and interpreted as evidence for full circular intermediates. Reads with one alignment segment mapping to the IS body and the other mapping to the adjacent chromosomal sequence were classified as *genome-head* (GH) or *tail-genome* (TG) junction reads, defining the corresponding insertion boundaries. IS110 candidates supported by at least one TH junction read were retained as high-confidence circularization events for downstream analyses.

### Local Assembly and IS110 Annotation

For IS110 candidates supported by circularization evidence, supporting long reads were extracted and assembled *de novo* using miniasm (v0.3-r179) [35] with overlap detection performed by minimap2 [36]. Consensus assemblies were polished using minipolish (v0.2.0) [65] to generate high-fidelity insertion sequences containing the complete IS body together with up to 80 bp of upstream and downstream flanking genomic sequence. When local assembly failed to converge, the longest supporting read was retained as the representative consensus sequence.

Open reading frames (ORFs) were predicted from each assembled insertion sequence using Prodigal (v2.6.3) [28] in metagenomic mode (`-p meta`) to enable robust gene prediction without requiring organism-specific training. Predicted protein sequences were subsequently searched against two profile Hidden Markov Models (HMMs), `DEDD Tnp IS110` (DEDD recombinase) and `Transposase 20` (Tnp20 transposase), formatted for HMMER3/f (v3.3) [19]. Searches were performed using HMMER (v3.4, `hmmsearch`) [19] or pyhmmer (v0.11.2) [33] with an *E*-value threshold of 1 × 10*^−^*^5^.

An insertion was classified as an IS110 element when a single predicted ORF produced significant matches to both HMM profiles or when the two domains were distributed across two adjacent ORFs separated by ≤ 300 bp, consistent with a split DEDD–Tnp20 architecture. Insertions matching only one of the two HMM profiles were excluded from downstream analyses.

### Detection of Partial Circular Intermediates

Partial circular intermediates were defined as circular recombination products derived from internal regions of an IS110 element rather than the complete insertion sequence. To detect these events, a non-redundant reference database of all identified IS bodies was constructed for each sample, and raw long reads were aligned using minimap2 [36] with the `-x map-ont` preset. Name-sorted BAM files were parsed using pysam [49] to identify split-read alignments consistent with partial circularization.

For reads containing multiple alignment segments on the same IS reference, alignment segments were ordered according to their query coordinates, and adjacent segment pairs were examined for a back-jump configuration in which the second alignment began upstream of the end of the preceding alignment on the reference sequence, consistent with circular junction formation. Candidate partial circles were required to span between 50 bp and 80% of the total IS length, thereby excluding full-length circular intermediates, and to contain at least 10 bp of aligned sequence on both sides of the predicted junction.

Breakpoint coordinates supported by independent reads were grouped using single-linkage clustering with a 5 bp distance tolerance. Clusters supported by at least two independent reads were retained as high-confidence partial circular intermediates for downstream analyses.

### Cargo Gene Annotation and Diversity Analysis

Accessory genes carried within IS110 candidate regions were annotated using a tiered functional annotation workflow combining gene prediction, specialized mobile-element databases, defense-system identification, and broad functional classification.

#### Coding Sequence Prediction

Coding sequences (CDSs) were predicted from nucleotide sequences derived from the empty-versus-filled boundary discovery pipeline after removal of flanking host genomic DNA using Prodigal (v2.6.3) in metagenomic mode (`-p meta`) [28]. To recover short, fragmented, or non-canonical open reading frames (ORFs) that may be missed by ab initio gene prediction, six-frame translations were additionally generated using EMBOSS `transeq` (v6.6.0.0) [51]. Supplementary ORFs were retained only when they were at least 100 amino acids long, did not overlap Prodigal-predicted CDSs, and received at least one downstream functional annotation. FASTA sequence identifiers were sanitized prior to downstream analyses to ensure compatibility across annotation software.

#### Functional Annotation

Predicted proteins and nucleotide sequences were annotated using complementary specialized databases to maximize functional coverage. Antimicrobial resistance (AMR), virulence, plasmid replicons, and metal/biocide resistance genes were identified using ABRICATE (v1.4.0) [53] with default thresholds (≥ 80% nucleotide identity and ≥ 80% coverage) against the CARD [1], ResFinder [6], NCBI Bacterial Antimicrobial Resistance Reference Gene Database [20], VFDB [37], PlasmidFinder [10], and BacMet2 [43] databases.

Anti-phage defense systems were identified from predicted proteins using DefenseFinder (v1.x) [63], incorporating its bundled defense-system profile HMM library [64], CasFinder profiles [14], and the MacSyFinder2 search framework [41]. Broader functional annotations were assigned using eggNOG-mapper (v2.1.13) [9] in DIAMOND search mode (`-m diamond`) [7] against the eggNOG database (v5.0.2) [26] to assign COG functional categories [22] and KEGG pathways [30].

#### Composition and Enrichment Analysis

Functional composition was calculated as the percentage of accessory CDSs assigned to each annotation category ((*n*_class_*/n*_total_) × 100%). Random-genome negative controls were generated from five independent sampling seeds, and the percentage of each functional category was summarized as the mean ± standard deviation across replicates. Functional enrichment within IS110 cargo was quantified as fold enrichment relative to the negative-control mean together with the corresponding *z*-score, calculated as

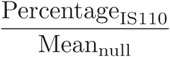

and

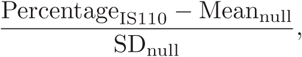

respectively. Functional categories absent from all negative-control replicates were reported as having undefined fold enrichment and *z*-scores.

### Cross-Genome Comparison of Large *IS110* Elements

#### Candidate Filtering and Clustering

To evaluate horizontal transfer of expanded *IS110* loci, we analyzed large *IS110* elements (≥ 5 kb, including the transposase and associated cargo) identified across complete genome assemblies. Candidate elements were required to encode an intact *IS110* transposase containing both the DEDD recombinase (`PF01548`) and Tnp20 (`PF02371`) domains within a single open reading frame (ORF). Redundant elements were clustered at 99% nucleotide identity using CD-HIT-EST, yielding 1,244 representative *IS110* sequences for downstream cross-genome analysis.

#### Database Alignment and Stringent Filtering

Representative *IS110* sequences were aligned against complete bacterial genome databases using Minimap2 (asm10 preset). Candidate transfer events were retained only when:

1. The aligned *IS110* sequence exhibited ≥ 99% nucleotide identity and ≥ 95% query coverage.
2. Both the source and target loci encoded intact two-domain *IS110* transposases.

This stringent sequence-level filter minimized the inclusion of fragmented or unrelated insertion sequences while restricting comparisons strictly to highly conserved *IS110* loci.

#### Flanking Genomic Context and Transfer Classification

To distinguish recent horizontal transfer from vertically inherited insertions, the genomic context surrounding each *IS110* element was compared using flanking chromosomal sequences. For every candidate pair, 5-kb regions immediately upstream and downstream of both *IS110* loci were extracted and aligned using Minimap2 through the `mappy` interface (with strand correction applied prior to comparing reverse-strand matches).

A flank was considered conserved when an alignment covered ≥ 10% of the 5-kb region with ≥ 90% nucleotide identity. Based on these conservation criteria, candidate transfers were classified into four categories:

- `MOVED:` Neither the upstream nor downstream flank showed detectable conservation.
- `SYNTENIC:` Both upstream and downstream flanks were conserved.
- `PARTIAL:` Only one flank (upstream or downstream) showed detectable conservation.
- `NO TGT FLANK:` Insufficient flanking sequence was available due to contig boundaries.

Only events classified as MOVED were retained as high-confidence horizontal transfer events.

#### Classification Rationale

The permissive 10% flank-coverage threshold was intentionally selected to maximize the detection of residual synteny. Under this criterion, any detectable conservation between flanking regions classified a locus as syntenic rather than horizontally transferred. Consequently, *IS110* loci classified as MOVED represent the most conservative set of transfer events, exhibiting highly conserved *IS110* bodies embedded within completely distinct genomic contexts.

### IS110 Lineage Detection

To reconstruct the progressive expansion of IS110 loci, we developed a computational workflow that identifies related structural variants occupying the same chromosomal insertion site across complete bacterial genome assemblies. For each insertion site anchored by a common pair of flanking sequences (*v*1_ref_id), all observed structural variants, including empty target sites (*V*_0_), canonical IS110 insertions (*v*_ref), deletion derivatives, and insertion derivatives, were retrieved from the genomic database. To reduce redundancy arising from minor assembly-length variation, variants were sorted by sequence length and collapsed into representative length clusters using a 50 bp threshold.

Evolutionary lineages were reconstructed by comparing all variants from the same insertion site using all-versus-all pairwise alignments generated with minimap2 (v2.30) [36] using the `asm10` preset and the `--eqx` option. Variants were assigned to the same lineage when the shorter sequence was contained within the longer sequence with at least 95% nucleotide identity and 95% query coverage. Variants that failed to satisfy these criteria initiated independent evolutionary lineages.

To remove highly truncated or degenerate insertion fragments, reconstructed lineages were further filtered by requiring the shortest sequence in each lineage to retain at least 95% of the canonical IS110 transposase coding sequence. This filtering step excluded pseudogene-like remnants while preserving complete IS110 expansion series. Final lineage structures were visualized using pyGenomeViz [54] together with matplotlib [27].

### Detection of Tandem Expansion within IS110 Loci

To identify tandem expansion events within IS110 loci, we combined self-alignment analysis with independent repeat-detection algorithms and randomized control datasets. Reference IS110 elements and their structural variants (≥ 1, 500 bp) were aligned against themselves using minimap2 (v2.30) [36] with main-diagonal masking (`-X`) to detect adjacent same-strand duplications. Candidate tandem repeats were required to exhibit at least 95% nucleotide identity, repeat units of at least 50 bp, and a minimum of two repeat copies. Overlapping harmonic repeat predictions were collapsed into a single representative array.

Candidate tandem arrays were independently validated using Tandem Repeats Finder (TRF v4.10.0rc2) with a minimum alignment score of 50 and maximum repeat period of 500 bp, together with ULTRA (v1.2.1), an HMM-based tandem repeat detection algorithm. Only tandem arrays supported by both methods were retained for downstream analyses.

To distinguish biologically enriched tandem expansions from background sequence composition, validated arrays were compared with two independent negative controls: (i) length-matched random genomic windows and (ii) IS110 sequences randomized using the Altschul–Erikson dinucleotide-preserving shuffling algorithm. Tandem repeat frequencies and array densities were compared using Fisher’s exact test, Mann–Whitney U tests, and 10,000-label permutation tests.

### Rearrangement Detection and Orthogonal Validation

Structural rearrangements associated with IS110 loci were identified by comparing the genomic positions and orientations of flanking anchor pairs between reference and target assemblies. Anchor pairs mapping to the same contig in the same orientation were classified as canonical loci. Candidate rearrangements were classified as inversions when anchor pairs mapped to the same contig but on opposite strands, translocations when the two anchors mapped to different contigs, and duplications when multiple high-confidence alignments were detected for either anchor. The 50 highest-confidence candidates from each rearrangement category for each species were selected for independent validation.

For each candidate event, local genomic regions spanning the reference IS110 locus (±30 kb) together with the corresponding target regions (±30 kb) were extracted and evaluated using two complementary whole-genome alignment approaches. MUMmer4 dnadiff (v4.0.1) [39] was used to identify localized inversions, insertions, and structural breakpoints from the generated `.qdiff` reports. ProgressiveMauve (v2.4.0) [17] was independently used to compare locally collinear blocks (LCBs), with inversions defined by reversed LCB orientation and translocations by changes in LCB order between the reference and target sequences. Because progressiveMauve does not explicitly resolve duplicated regions, duplication events were evaluated using dnadiff alone.

To minimize false-positive translocation calls arising from fragmented draft assemblies, candidate translocations with either anchor located within 5 kb of a contig terminus were classified as unresolved contig-break artifacts and excluded from further analysis. Final rearrangement calls required concordant support from both dnadiff and progressiveMauve for inversions and translocations, whereas duplication events required confirmation by dnadiff alone. Validation of the sampled candidate set confirmed 229 inversions (65% of 350 candidates) and 258 duplications (73% of 351 candidates).

### Cross-Organism Transfer of Large IS110 Elements

To identify recent horizontal transfer of large IS110 elements, all IS110 loci ≥ 5 kb identified from the empty-versus-filled boundary pipeline were collected from ten bacterial species. Whole-element nucleotide sequences were dereplicated using CD-HIT-EST (v4.8.1) [21] with the parameters -c 0.99 -aS 0.95, yielding 1,244 representative IS110 sequences. Representative elements were required to encode an IS110 transposase containing both conserved Pfam domains (PF01548 and PF02371).

Representative IS110 sequences were aligned against the complete IS110 datasets of all species using minimap2 (v2.30) [36] with the `-x asm10 -c --eqx` options and pre-built `.mmi` reference indexes. Cross-species matches were retained when the source and target species differed, alignment coverage was at least 30% of the query sequence, and weighted nucleotide identity was at least 95%. A high-confidence subset representing recent transfer events additionally required at least 99% nucleotide identity and 95% query coverage.

To distinguish transfer of complete IS110 elements from sequence similarity to non-IS genomic regions, candidate alignments were intersected with the independently generated HMM-based IS110 annotation catalogue, requiring co-localization with a predicted IS110 transposase. Taxonomic relationships between source and target genomes were recorded at the family, class, and phylum levels.

To reduce false-positive horizontal transfer events arising from contaminated or misannotated genome assemblies, cross-family matches were further evaluated according to their distribution across target assemblies. Candidate transfers were classified as `SUSPECT CONTAMINATION` when at least 50% of all hits were concentrated within three or fewer target assemblies, `ROBUST HGT` when at least five independent target assemblies each contributed less than 40% of the total hits, and `MODERATE` for intermediate cases.

Each retained source–target genome comparison was counted as one pairwise IS110 match. Because multiple assemblies can represent closely related bacterial isolates, these pairwise counts were used to quantify the density of transfer relationships and were not interpreted as counts of independent horizontal-transfer events.

### Identification of Plasmid Capture by Large *IS110* Elements

To identify large *IS110* elements carrying integrated plasmid sequences, all *IS110* candidate bodies ≥ 10 kb were screened for plasmid replication and transfer signatures. Open reading frames (ORFs) were predicted using Prodigal (v2.6.3) in metagenomic mode (`-p meta`) [28]. Replication initiation (*rep*) proteins were identified using MOB-suite (v3.1.9; `mob typer`) [52]. Origin-of-transfer (*oriT* ) sites were detected using blastn searches against the oriTDB2 database with an *E*-value threshold of 1 × 10*^−^*^10^ and a minimum nucleotide identity of 80%. Vegetative origins of replication (*oriV* ) were identified using blastn against a combined DoriC–PLSDB reference database with a minimum nucleotide identity of 90% and an alignment length of at least 200 bp.

Candidate plasmid-capture events were defined as *IS110* elements simultaneously encoding all three plasmid-associated signatures (*rep*, *oriT*, and *oriV* ). To restrict downstream analyses to chromosomally integrated plasmid fragments and exclude plasmid assemblies, candidates were retained only when located on contigs at least 1 Mb in length.

### Genomic DNA Extraction, Circular DNA Enrichment, and Rolling Circle Amplification

Wild-type *Escherichia coli* strains (s186, s189, and s193) were streaked onto antibiotic-free Luria–Bertani (LB) agar plates and incubated at 37 °C. Single colonies were inoculated into LB broth and cultured for 6–8 h at 37 °C. Cells were harvested by centrifugation (16, 000 × *g*, 1 min), lysed in Tissue and Cell Lysis Solution containing Proteinase K at 65 °C for 15 min, and treated with 1 *μ*L RNase A (5 mg/mL) at 37 °C for 30 min to remove RNA. Proteins were precipitated using MPC Protein Precipitation Reagent (16, 000 × *g*, 4 °C, 10 min), and total nucleic acids were recovered by isopropanol precipitation, washed twice with 70% ethanol, and resuspended in nuclease-free water overnight at 4 °C.

To enrich circular DNA species, genomic DNA was mechanically sheared by repeated vortexing and pipetting before overnight digestion with Plasmid-Safe ATP-Dependent DNase at 37 °C. The enzyme was heat-inactivated at 70 °C for 30 min, and depletion of linear genomic DNA was verified by PCR amplification of the 16S rRNA gene followed by electrophoresis on a 2% agarose gel.

Circular DNA was subsequently amplified by rolling circle amplification (RCA) using *φ*29 DNA polymerase and target-specific phosphorothioate-protected primers at 30 °C for 12 h. Following heat inactivation at 65 °C for 10 min, amplified products were diluted and subjected to Oxford Nanopore long-read sequencing to identify circular junction reads.

### Nanopore RCA Read Processing, Transposase Domain Typing, and Cargo Circle Profiling

Nanopore rolling circle amplification (RCA) reads generated from two sBX0186 replicates (barcodes 80 and 81) were processed using TideHunter [23] to generate high-accuracy consensus monomer sequences.

Consensus monomers were converted to FASTA format and aligned to the sBX0186 reference genome using minimap2 with the `map-ont` preset [36].

To identify authentic *IS110* elements, predicted reference coding sequence (CDS) proteins were screened using `hmmsearch` [19] against the Pfam Tnp 20 (PF02371) and DEDD (PF01548) profile HMMs. Candidate transposases were required to contain both conserved domains for classification as bona fide *IS110* transposases.

Primary and supplementary monomer alignments were integrated to classify *IS110* CDS coverage as FULL (complete CDS coverage), `PARTIAL`, or `ABSENT`. For each locus, the unique full-length monomer spanning the complete *IS110* transposase CDS was selected as a local reference sequence, and all remaining monomers were remapped against this reference to identify shorter circular intermediates and IS-only circular species originating from the same locus.

To characterize excision boundaries, the first and last 7 bp of each linearized consensus monomer were compared pairwise using a Hamming distance threshold of at most one mismatch (≥ 6*/*7 matching nucleotides) to identify shared circularization junctions.

## Data availability

The complete dataset, including the catalogue of detected and verified IS110-mediated genomic rearrangements, is openly available on Hugging Face at https://huggingface.co/datasets/hukuang/IS110_repo. The source code, custom scripts, and software pipeline for identification and validation are hosted on GitHub at https://github.com/KuangHu/IS110_fragment.

## Software and Database Inventory

To ensure reproducibility, all bioinformatics tools, reference databases, and execution libraries utilized across the computational pipelines are systematically cataloged below (Table 1).

**Table 1.**
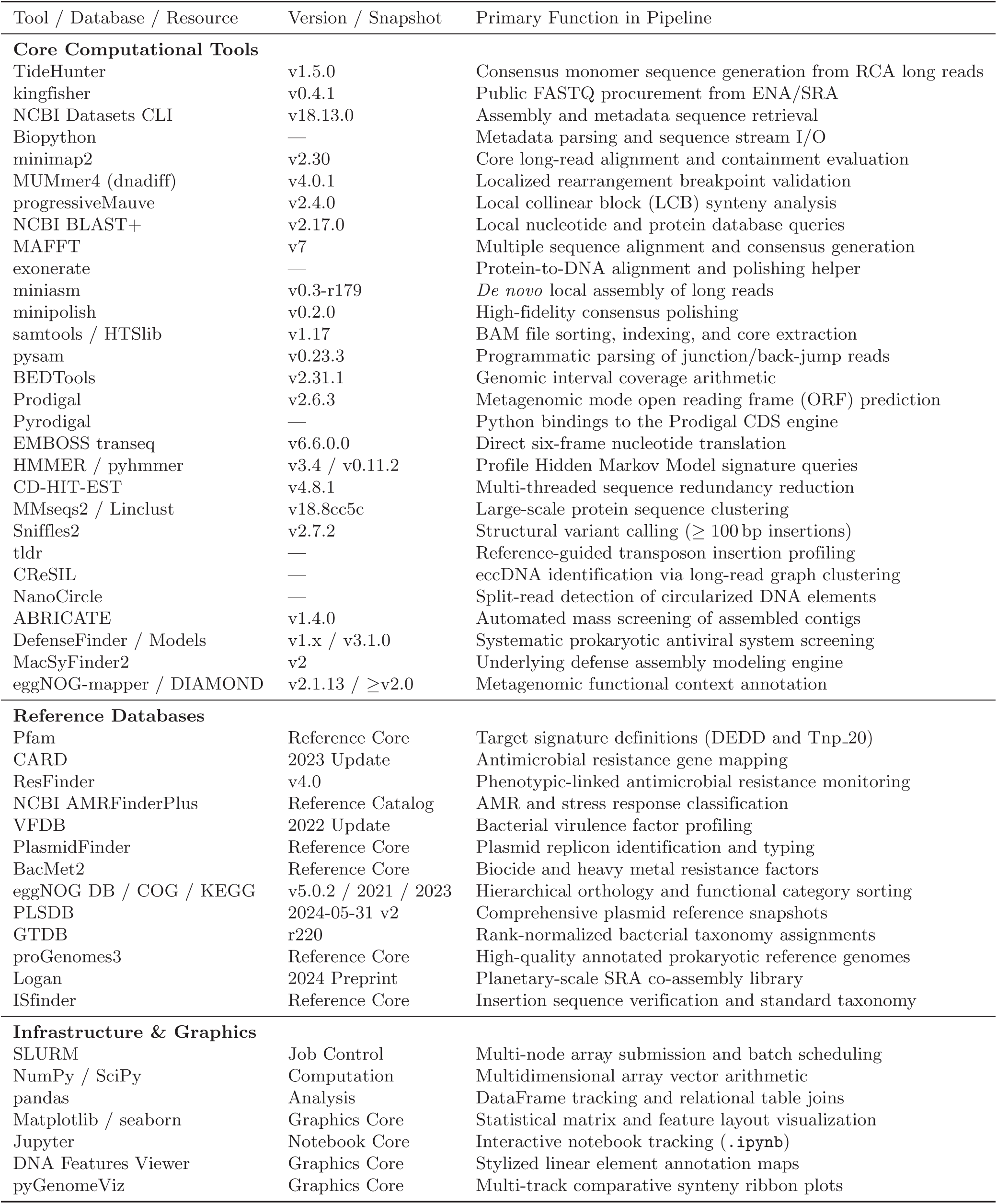
Comprehensive inventory of bioinformatics software, databases, and visualization libraries utilized in this study.

## Acknowledgements

We express our gratitude to Brady Cress for generously offering useful insights into the IS110 mechanism. The authors utilized Claude (Anthropic) for coding assistance and editing the text. However, all code, analyses, and manuscript content were checked and verified by the authors, who bear full responsibility for the work.

We’d like to thank our funders:

- **JBEI:** This material is based upon work at the Joint BioEnergy Institute (JBEI) supported by the U.S. Department of Energy, Office of Science, Biological and Environmental Research Program under contract DE-AC02-05CH11231 with Lawrence Berkeley National Laboratory.
- **The Audacious Project:** This work was supported in part by Lyda Hill Philanthropies, Acton Family Giving, the Valhalla Foundation, Hastings/Quillin Fund—an advised fund of the Silicon Valley Community Foundation, the CH Foundation, Laura and Gary Lauder and Family, the Sea Grape Foundation, the Emerson Collective, Mike Schroepfer and Erin Hoffman Family Fund—an advised fund of Silicon Valley Community Foundation, and the Anne Wojcicki Foundation through The Audacious Project at the Innovative Genomics Institute.
- **Leona M. and Harry B. Helmsley Charitable Trust:** This work was funded in part by grant [G-2302-06692] from The Leona M. and Harry B. Helmsley Charitable Trust.
- **Shurl and Kay Curci Foundation:** This work was supported by a Research Award from the Shurl and Kay Curci Foundation (https://curcifoundation.org) to the Innovative Genomics Institute Genomic Tool Discovery Program at UC Berkeley, awarded to B.E.R.
- **Innovative Genomics Institute:** This work was supported in part by the Innovative Genomics Institute.
- **m-CAFEs:** This work was supported by m-CAFEs Microbial Community Analysis and Functional Evaluation in Soils, a Science Focus Area led by Lawrence Berkeley National Laboratory based upon work supported by the US Department of Energy, Office of Science, Office of Biological and Environmental Research under contract number DE-AC02-05CH11231.

