## Supplementary Figures for "Canonically minimal RNA-guided insertion sequences expand into large elements that disseminate antimicrobial resistance"

Kuang Hu, et al.

August 21, 2026

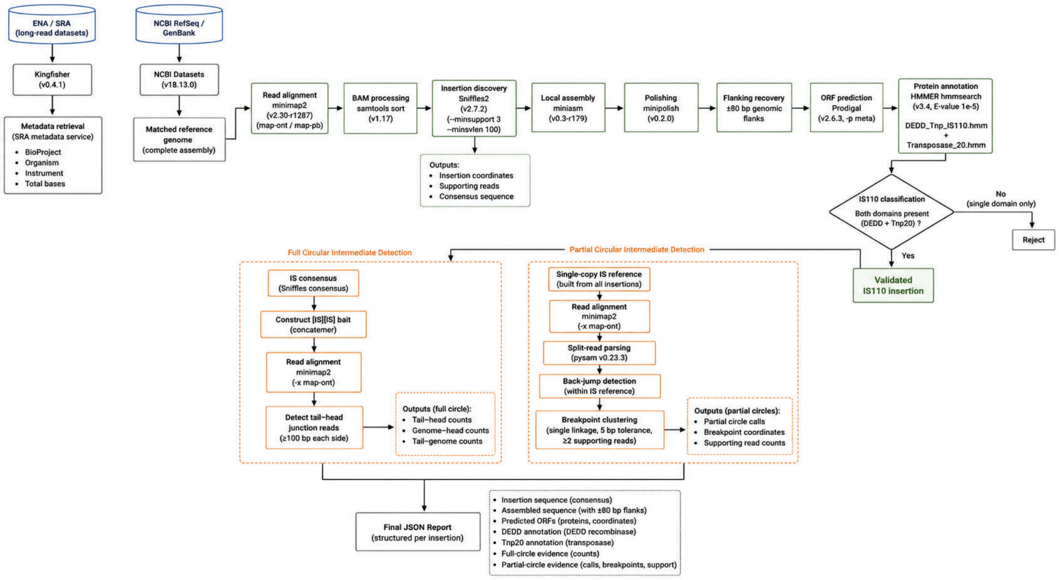


Figure S1: **Computational pipeline for IS110 discovery and circular intermediate detection from long-read sequencing.** Schematic overview of the workflow for IS110 identification, annotation, and circular intermediate detection using long-read sequencing data. Results from all analysis steps were integrated into a final report containing sequence and evidence summaries.

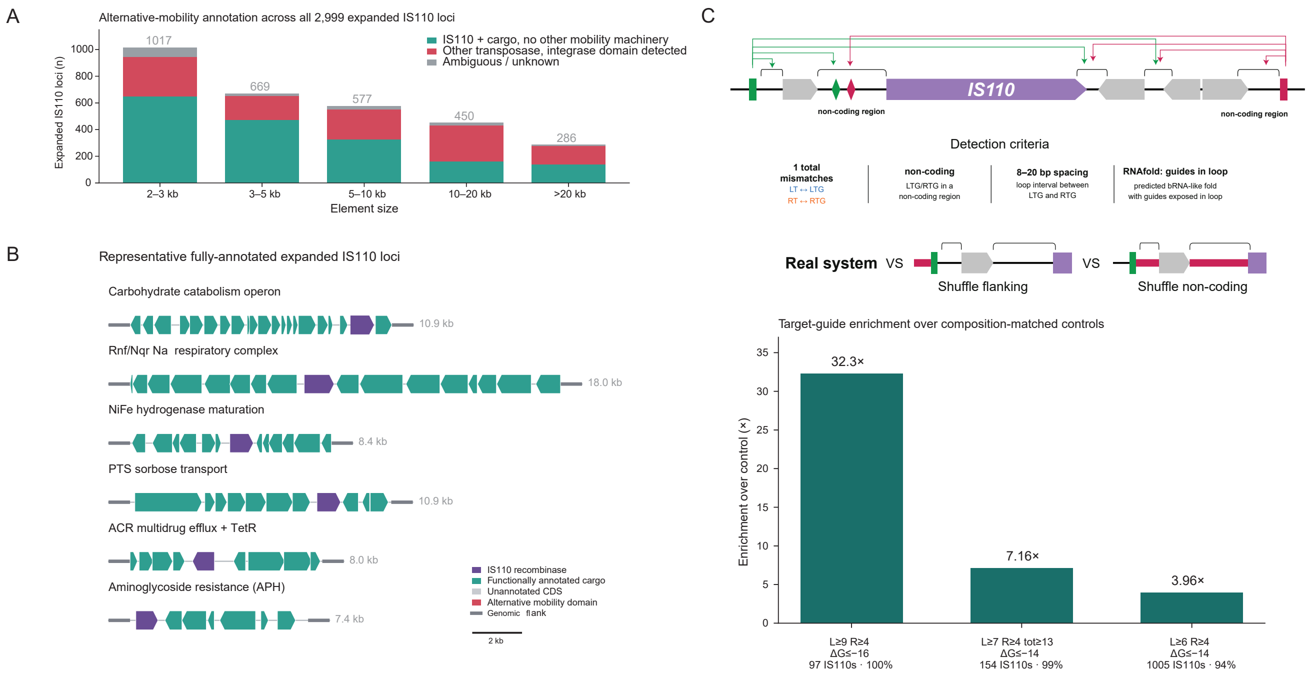


Figure S2: **Expanded IS110 loci retain IS110-specific architecture and contain predicted bridge-RNA target-guide signatures.** **A.** Alternative-mobility annotation of all 2,999 IS110 elements >2 kb, stratified by element length. Elements were classified as IS110 plus cargo with no detectable alternative transposase or integrase, containing an additional transposase/integrase domain, or having an ambiguous or incompletely annotated gene complement. Numbers above bars indicate the total number of elements per size bin. Overall, 1,474 cargo-bearing expanded IS110 loci lacked a detectable alternative transposase or integrase. **B.** Gene maps of six representative expanded IS110 loci in which all non-IS110 CDSs received Pfam-A functional annotations. Genes are shown to scale from Prodigal-predicted coordinates, with arrowheads indicating strand orientation. No additional transposase or integrase domain was detected in the loci shown, which carry representative metabolic, respiratory, transport, and antimicrobial-resistance cargo. Scale bar, 2 kb. **C.** Detection of predicted bridge-RNA target-guide signatures within IS110 bodies. *Top:* candidate LTG and RTG sequences in non-coding regions of the IS110 body were matched to the corresponding left and right target flanks (LT and RT) of the pre-insertion site and filtered by sequence complementarity, guide spacing, and RNAfold-predicted loop exposure. *Middle:* authentic body–flank pairings were compared with composition-matched controls generated by shuffling either the target-flanking sequence or the non-coding IS110 sequence. *Bottom:* enrichment relative to the mean of the two controls under TIGHT ($L \geq 9$ nt, $R \geq 4$ nt, $\Delta G \leq -16$ kcal mol$^{-1}$; 97 elements; 32.3×), MID ($L \geq 7$ nt, $R \geq 4$ nt, combined guide length $\geq 13$ nt, $\Delta G \leq -10$ kcal mol$^{-1}$; 154 elements; 7.16×), and WIDE ($L \geq 6$ nt, $R \geq 4$ nt, $\Delta G \leq -14$ kcal mol$^{-1}$; 1,005 elements; 3.96×) criteria. Enrichment was calculated using a +1 pseudocount in the control denominator.

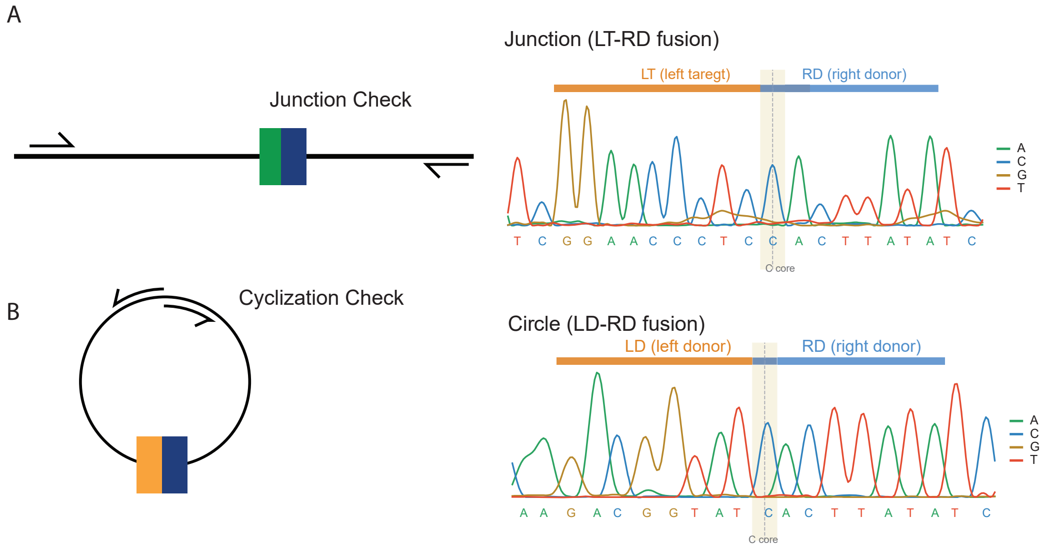


Figure S3: **Sanger validation of IS110 circularization and insertion junctions. A.** Representative Sanger sequencing chromatogram confirming the expected LT–RD junction generated by IS110-mediated insertion. Junction positions are indicated by dashed lines. **B.** Representative Sanger sequencing chromatogram confirming the expected LD–RD junction generated by IS110 circularization. Original .ab1 files are provided as Supplementary Source Data.

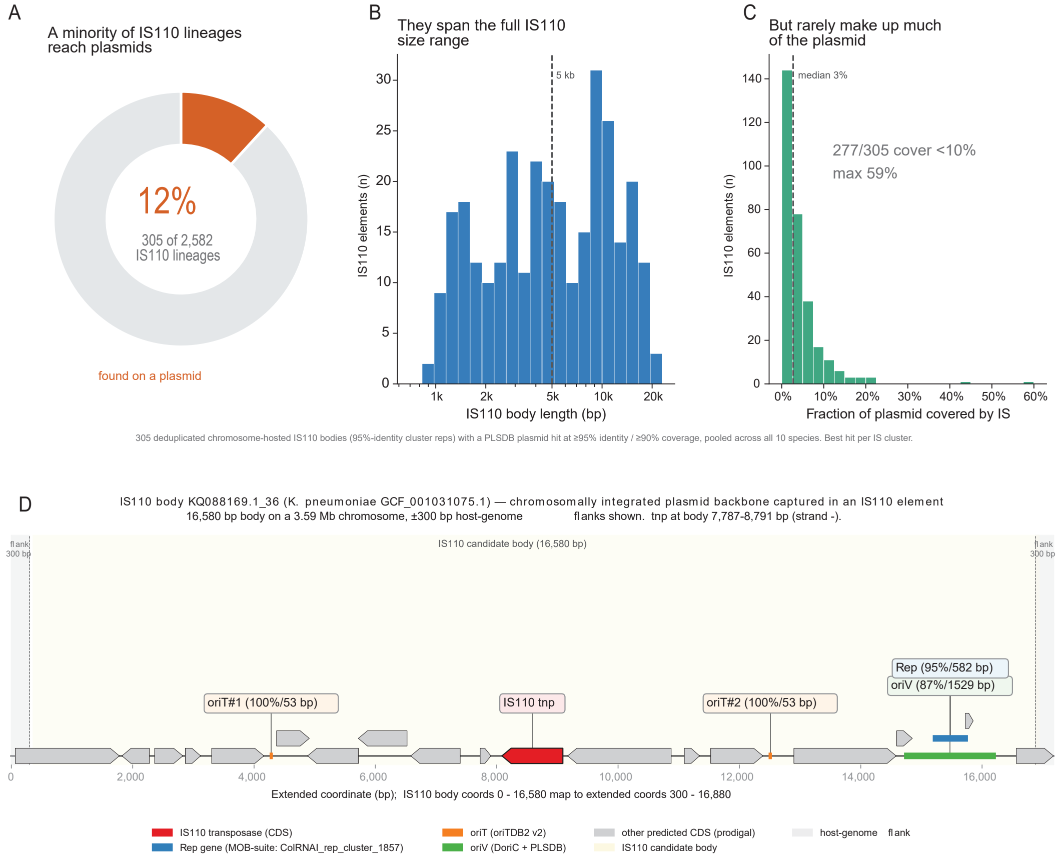


Figure S4: **Chromosome–plasmid sharing of IS110 elements and capture of plasmid replication modules.** Chromosomal IS110 elements were aligned against the PLSDB plasmid database (minimap2, asm10). Matches with $\geq$95% nucleotide identity and $\geq$90% query coverage were retained, with one best hit per 95%-identity IS110 cluster, yielding 305 deduplicated chromosome-to-plasmid "shuttle" events. Analyses are pooled across all ten bacterial species. **A.** Of 2,582 unique IS110 lineages reconstructed from complete genome assemblies (95%-identity cluster representatives), 305 (12%) have a high-confidence plasmid match, indicating that nearly identical IS110 elements occur in both chromosomal and plasmid contexts. **B.** These shared elements span the full observed IS110 size range (median 4,901 bp; 151/305 $\geq$5 kb; dashed line, 5 kb; x-axis on a logarithmic scale). **C.** The IS110 body typically comprises only a small fraction of its host plasmid (median 3%; 277/305 <10%; maximum 59%; dashed line, median), indicating that transferred elements are embedded within larger plasmid backbones rather than constituting complete plasmids. Data underlying this figure are provided in `panel_shuttle_portion_source_data.csv`. **D.** Organization of a 16.6-kb chromosomal IS110 locus (`KQ088169.1_36`) from *Klebsiella pneumoniae*. The element contains an IS110 transposase together with a ColRNAI-family *rep* gene, one predicted *oriV*, and two predicted *oriT* sites, representing a captured plasmid replication and transfer module. Gray arrows denote predicted CDSs; 300-bp chromosomal flanks are shown.

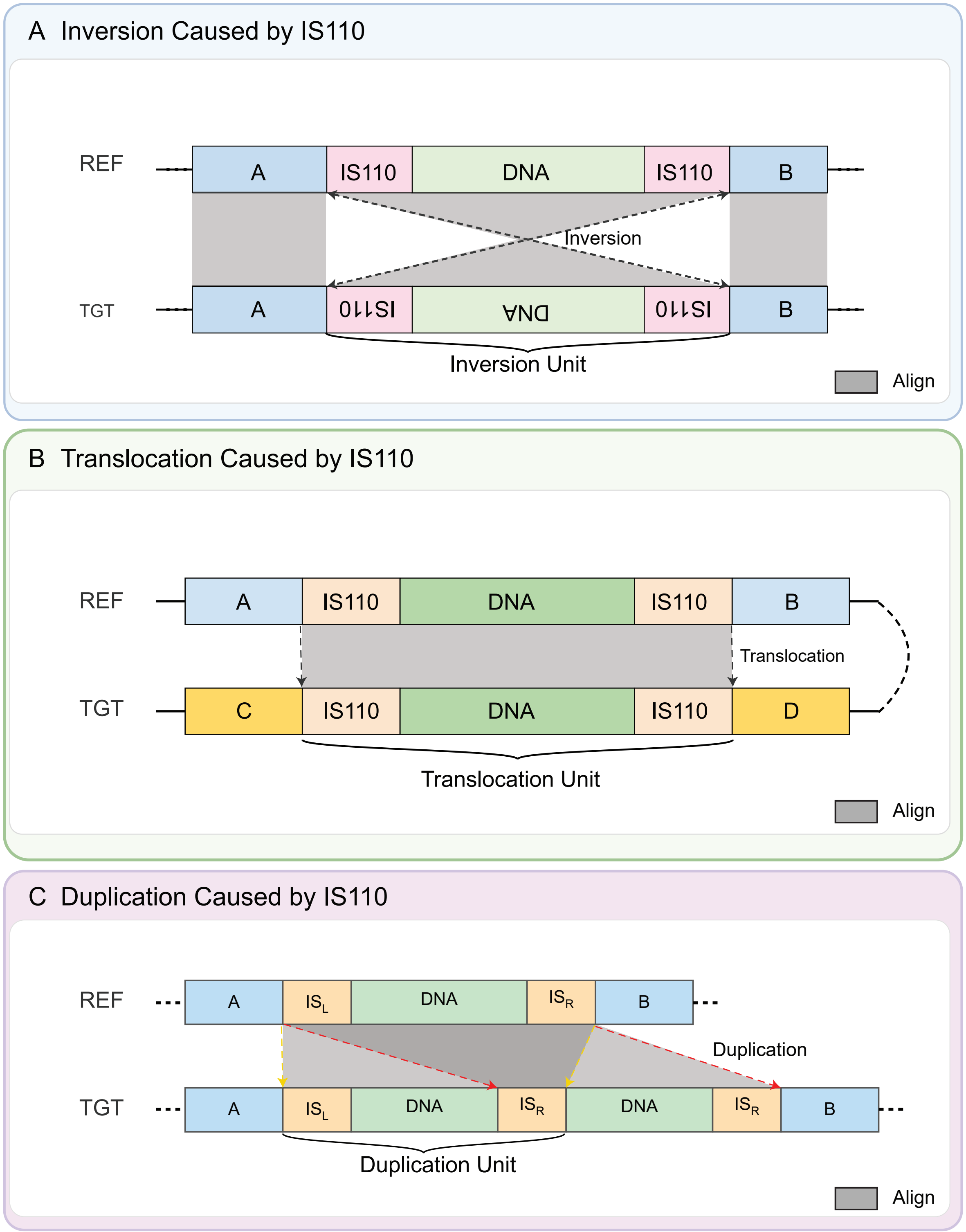


Figure S5: **Molecular mechanisms and canonical structural variations mediated by IS110 composite transposons. A.** Genomic inversion spanning an IS110-flanked cargo segment, characterized by syntenic forward alignments of outer flanks A and B, alongside a complete reverse-complement rearrangement of the internal inversion unit. **B.** Inter-contig translocation driven by composite transposon movement, validated by a distinct sequence context shift where reference flanking regions A and B fail to match the novel target insertion sites C and D. **C.** Multi-copy tandem duplication and sequence expansion, where reference locus containing a single cargo segment maps to multiple duplicated target alignment blocks separated by functional inter-copy insertion sequences.

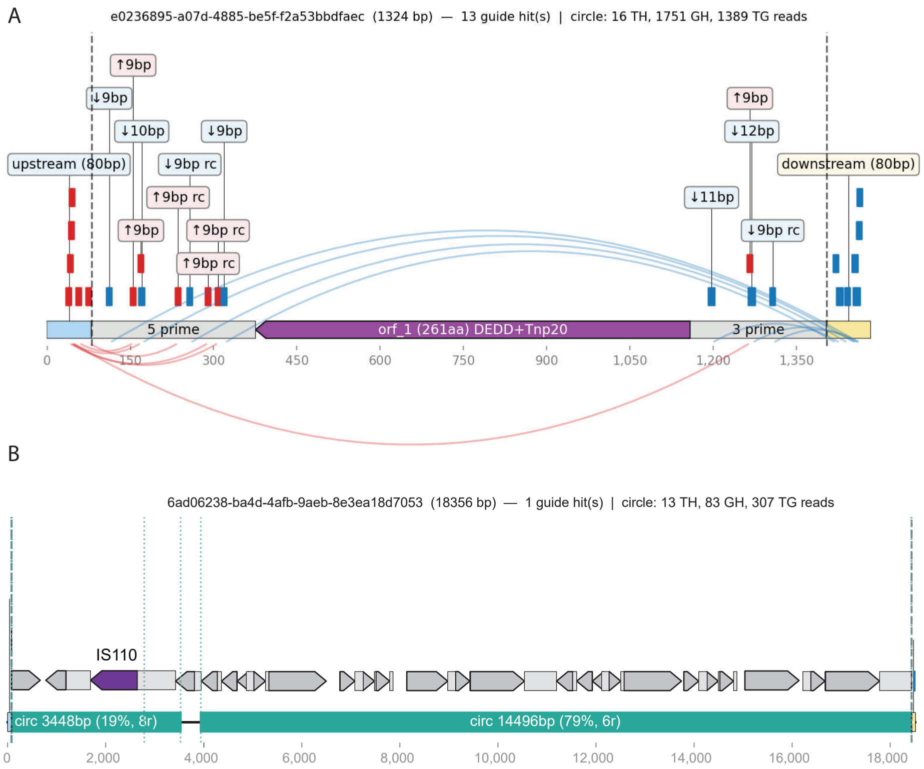


Figure S6: **Representative read-mapping views supporting active IS110 recombination. A.** Canonical IS110 locus showing guide annotations together with circular intermediate and junction-read support. **B.** Expanded IS110 locus showing multiple partial circular intermediates but no evidence of whole-element circularization.

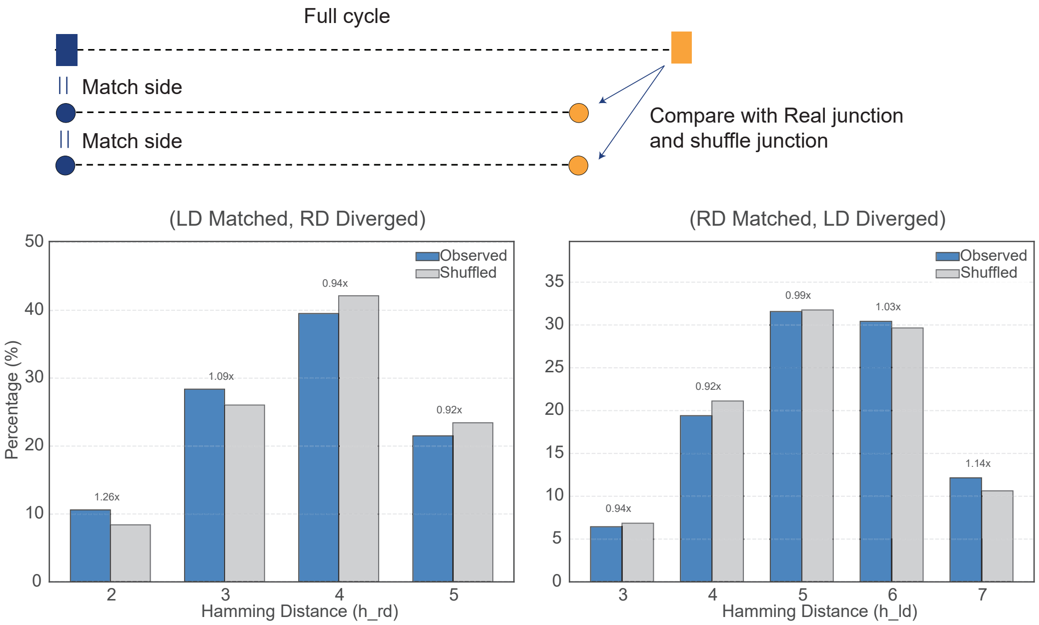


Figure S7: **Hamming Distance Distribution of Diverged Junctions in Matched Circle Intermediates.** Comparison of observed and shuffled junction sequences to assess sequence similarity at the unmatched donor boundary of partial circular intermediates. The bar charts show the distribution of Hamming distances at the divergent junction side for LD-only and RD-only partial-circle junctions, in which the opposite donor boundary retains a canonical match. Observed sequences were compared with shuffled junction controls generated from the corresponding loci. Fold-change values indicate the ratio of observed to shuffled frequencies at each Hamming distance. The close agreement between observed and shuffled distributions indicates that the unmatched boundary shows no greater similarity to the corresponding canonical donor sequence than expected by chance.

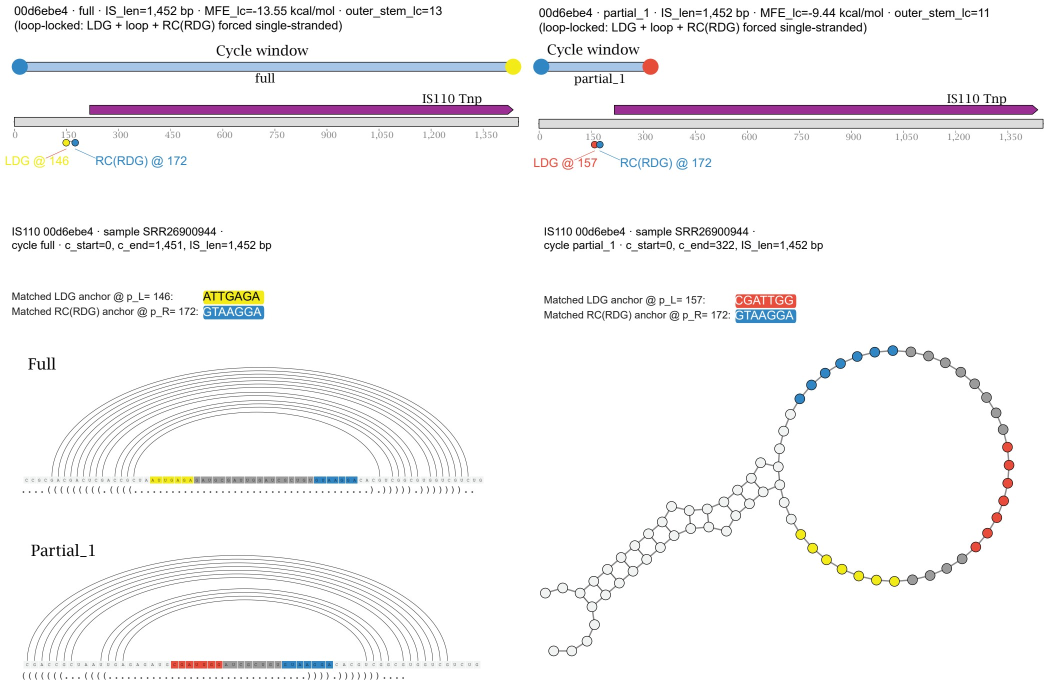


Figure S8: **Alternative positioning of donor-recognition sequences within the bRNA loop may explain an apparently divergent partial-circle junction.** Comparison of a full-length and a partial circular intermediate from the same 1,452-bp IS110 element. In the full circle, the LDG and RC(RDG), located at positions 146 and 172, respectively, perfectly match the corresponding donor boundaries and are separated by 19 nt. In the partial circle, an alternative LDG-like sequence located 11 nt downstream, at position 157, matches the left donor boundary with one mismatch, whereas the same RC(RDG) retains a perfect match to the right donor boundary, resulting in an 8-nt anchor separation. Predicted bRNA structures place both LDG candidates and the RC(RDG) within the same exposed loop, suggesting that nearby loop sequences may function as alternative donor-recognition segments.

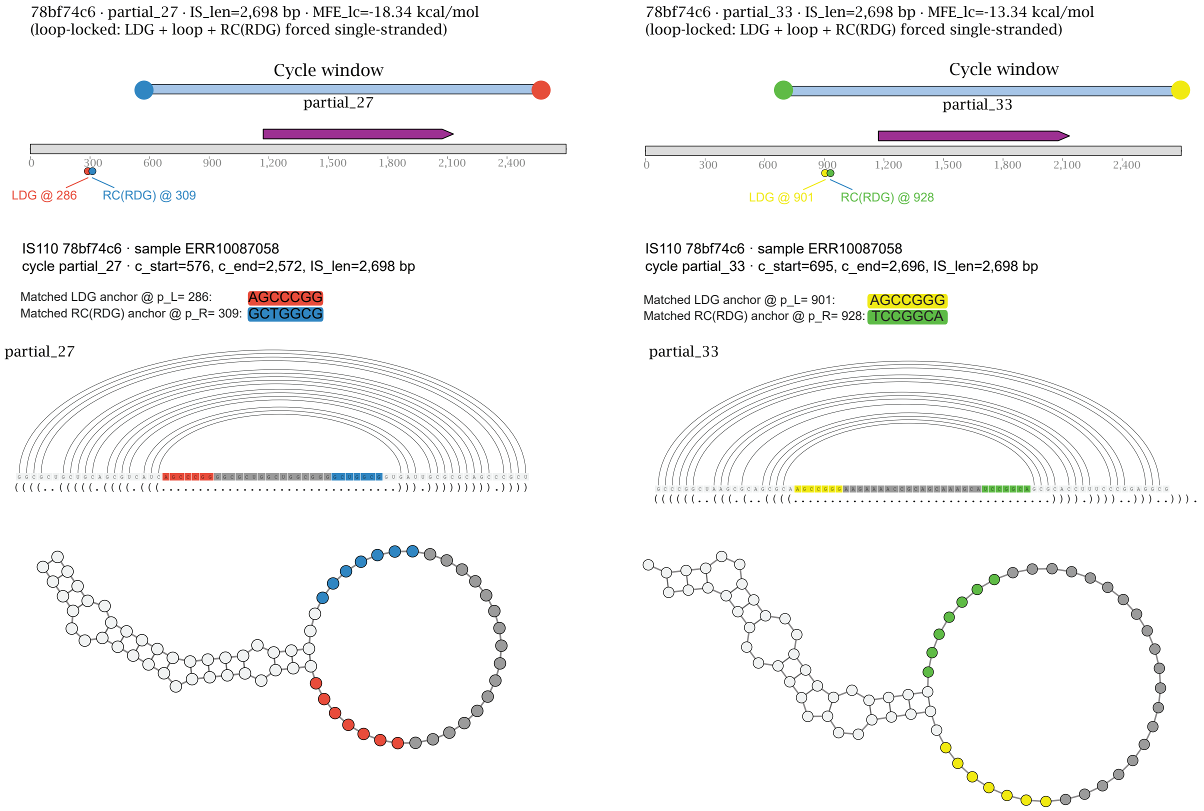


Figure S9: **Distinct predicted donor-binding loops are associated with different partial circularization events within the same IS110 element.** Two partial circular intermediates detected from the same 2,698-bp IS110 element were associated with separate donor-guide regions. Partial circle 27 was compatible with an LDG–RC(RDG) pair located at positions 286 and 309, whereas partial circle 33 was associated with a second pair located at positions 901 and 928. In both cases, the predicted guide sequences showed near-complementarity to the corresponding left and right donor boundaries and were positioned within exposed RNA loop structures. These observations suggest that a single IS110 locus may encode multiple donor-binding loops that recognize distinct donor-boundary pairs. Apparent divergence of partial-circle junctions may therefore reflect assignment to a non-cognate donor-binding loop rather than guide-independent recombination.

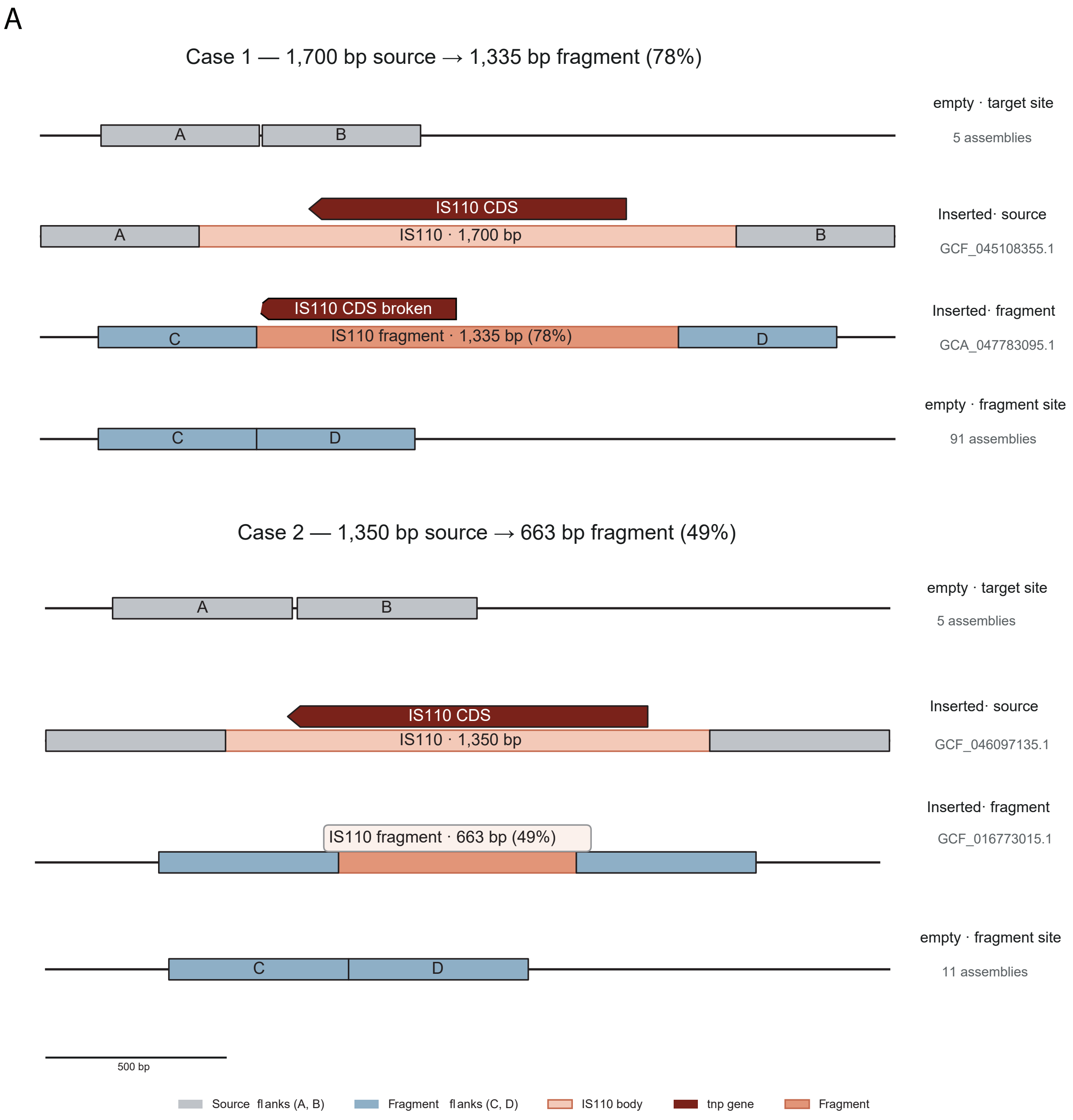


Figure S10: **Comparative genomic evidence for insertion of partial IS110 intermediates.** Two representative cases showing a full-length IS110 element at the donor locus (A–IS110–B) and a highly similar partial IS110 fragment ($>$ 99% nucleotide identity) inserted at an independent genomic locus (C–fragment–D), together with the corresponding empty donor and target sites. These examples support genomic reintegration of partial IS110 intermediates.
